# CMTM6-Silencing Microbial Immunotherapy Reprograms PDAC Tumors and Restores T-cell Function

**DOI:** 10.64898/2026.01.26.701790

**Authors:** CY Chabu, R Kazmierczak, M Hasani, N Patterson, Q Wang, L Canti, M.Z Tesfay, A Cios, B Dhagat, M.Q Pastor, C De la Nuez, T Verburg, J Moyer, K Gunter, M Mwanza, O Moaven, G Li, P de Figueiredo, B.M Nagalo

## Abstract

Despite recent advances in immunotherapy for advanced malignancies, Pancreatic ductal adenocarcinoma (PDAC) remains largely refractory to current immunotherapy due to dense fibrosis, limited antigen presentation, and myeloid-driven immune suppression. Here we report the tumor-targeting, immune remodeling, and safety profiles of the attenuated Salmonella enterica serovar Typhimurium strain CRC2631, and of iSTORM, a next-generation derivative engineered for tumor-localized CMTM6 silencing. CRC2631 preferentially colonizes orthotopic and genetically engineered PDAC tumors, with enrichment in primary lesions and metastases. Tumor-localized CRC2631 induces chemokine and adhesion programs consistent with leukocyte recruitment, increases intratumoral activated T-cell fractions, and triggers transcriptional signatures aligned with innate sensing, interferon signaling, antigen-processing and presentation, and apoptosis programs.

iSTORM extends this platform by delivering CMTM6-targeting shRNA to modulate a PD-L1-stabilizing, myeloid-associated immune-evasion programs within tumor-colonized tissue. Compared with CRC2631, iSTORM increases intratumoral CD8+ T cells, shifts T-cell state toward activation with reduced exhaustion-prone features, strengthens antigen-presentation programs, and achieves deeper tumor control. A lyophilized formulation preserves immune remodeling while improving deployability. Mechanistically, glycan arrays and functional studies support mannose-rich glycan-guided tumor engagement. iSTORM toxicity studies, including systemic cytokine, hematologic, blood chemistry, and lethality demonstrate a favorable safety profile.

Collectively, these findings establish iSTORM as a safe, programmable, CMTM6-silencing microbial immunotherapy platform that selectively targets and penetrate PDAC tumors to unleash anti-tumor immune activities.

**What is already known on this topic:** PDAC is highly resistant to immune checkpoint blockade because dense stroma and myeloid-dominated suppression prevent effective T-cell infiltration; attenuated *Salmonella* strains can selectively colonize tumors but first-generation agents showed limited efficacy and safety concerns.

**What this study adds:** This study defines CRC2631/iSTORM as a tumor-selective microbial immunotherapy that exploits surface-exposed, mannose-rich N-glycans to colonize PDAC, delivers CMTM6 silencing, and restores CD8^+^ T-cell activation and tumor control in models resistant to PD-1 blockade immunotherapy.

**How this study might affect research, practice or policy:** These findings provide a mechanistic blueprint for glycan-guided, CMTM6-targeted bacterial “living drugs,” support rational combination strategies for deepening therapeutic effect, and establish a lyophilized, biocontained platform that could be developed into scalable microbial immunotherapies for PDAC and other immunologically cold solid tumors.

## INTRODUCTION

Pancreatic ductal adenocarcinoma (PDAC) remains among the most lethal malignancies, with over 70% of patients presenting with metastatic disease and deriving minimal benefit from existing systemic therapies^1^. Even aggressive chemotherapy regimens offer limited durability, and most patients progress rapidly due to innate and acquired resistance mechanisms^2–5^. Despite substantial advances in cancer immunotherapy, immune checkpoint inhibitors, personalized neoantigen vaccines, and adoptive T-cell therapies have demonstrated minimal efficacy in PDAC outside of rare genomic subtypes (e.g., MSI-H or POLE-mutant tumors)^6^. These poor outcomes are due to the fibrous and immunosuppressive PDAC milieu.

Cancer-associated fibroblasts deposit dense extracellular matrix that compresses vasculature and restricts drug access^7–9^. In parallel, PDAC actively disables antitumor immunity by recruiting myeloid-derived suppressor cells (MDSCs) and suppressive macrophages (Tumor associated macrophages/TAMs) that block T-cell priming and function^10,11^. Further, cytokine-driven PD-L1 stabilization allows tumors to extinguish cytotoxic activity^12,13^. Strategies for overcoming PDAC physical and immune barriers are urgently needed.

Recent studies have identified CMTM6 as a central orchestrator of immune resistance in solid tumors, including PDAC^14–23^. CMTM6 is a MARVEL family transmembrane protein that not only stabilizes PD-L1 but also promotes MDSC recruitment, Treg activity, and ERK1/2-dependent polarization of macrophages into an immunosuppressive state^14–18,22^. High CMTM6 expression correlates with poor prognosis in PDAC and multiple solid tumors^18–20^; however in spite of its clear therapeutic relevance, no CMTM6-targeting agents currently exist, leaving a major immunologic opportunity unaddressed.

Microbial immunotherapy provides a compelling strategy to therapeutically exploit this vulnerability. Tumor-colonizing bacteria can selectively penetrate hypoxic, fibrotic tumor regions that are largely inaccessible to conventional biologics, while enabling localized and programmable delivery of immunomodulatory payloads^24–29^. Early efforts using attenuated Salmonella strains, such as VNP20009, demonstrated tumor tropism in murine models^30–32^. However, VNP20009 showed toxicity, immune clearance, and limited tumor control in clinical studies^24,33^. In contrast, advances in bacterial attenuation and metabolic engineering have substantially improved safety, tumor residency, and therapeutic precision^34,35^.

Several engineered *Salmonella* and *Listeria* strains have reached Phase I/II clinical evaluation stages (NCT03762291, NCT03190265, NCT03006302), and a genetically attenuated IL-2-expressing *Salmonella* (Saltikva) recently received FDA fast-track designation for metastatic PDAC (NCT04589234). These advances underscore the notion that bacterial platforms can be safe, deliver payloads directly within tumors, and elicit immune responses where that conventional therapies fail.

CRC2631 represents a distinct lineage within this therapeutic class. Developed from a genetically diverse LT2 population that evolved under nutrient restriction over four decades^36,37^, CRC2631 was attenuated through rfaH deletion and rendered auxotrophic for aromatic amino acids and thymine (aroA, thyA)^35,38–40^. These auxotrophies bias the strain toward tumors enriched in aromatic metabolites such as tryptophan and its immunosuppressive catabolites, a metabolic hallmark of PDAC and cold TMEs^41–43^, enhancing CRC2631 preferential growth in PDAC biochemical ecosystem.

We extended this platform by engineering iSTORM, the first microbial therapeutic designed to penetrate tumors and block CMTM6 directly within the TME, thereby destabilizing PD-L1 and dismantling CMTM6-driven myeloid suppression^14–18^. This dual mechanism (tumor-selective colonization and in situ immune reprogramming) addresses the two most entrenched barriers in PDAC: stromal exclusion and immunosuppression.

We demonstrate that CRC2631 penetrate tumor cores and overcomes checkpoint-refractory immunity, although suppressive myeloid activity emerges and coincides with limited effector CD8 T cell activity. iSTORM depletes CMTM6, partly overcomes myeloid recruitment and immunosuppression, and stimulate CD8 T-cells more robustly than CRC2631. Further, a lyophilized formulation activates systemic antitumor immunity and produces dose-dependent tumor suppression. These studies establish a readily scalable new platform for microbial immunotherapy in PDAC and set the stage for combination approaches aimed at deepening iSTORM therapeutic effects.

## RESULTS

### CRC2631 safely accumulates in primary PDAC tumors and metastatic sites

CRC2631 is a genetically attenuated *Salmonella enterica* strain previously shown to colonize tumors without overt toxicity in a prostate cancer model^35^. We sought to establish the extent to which this microbial platform can be translated to PDAC. We assessed CRC2631’s ability to preferentially colonize PDAC tissues in *in-vivo* biodistribution studies. CRC2631 was engineered to constitutively express iRFP720 and a chloramphenicol resistance cassette^35^. iRFP720 enables tissue near-infrared imaging, while chloramphenicol selection allows selective enumeration of viable CRC2631 from tissues, critical for distinguishing active tumor colonization from background flora.

Fluorescence IVIS imaging in the orthotopic Panc02H7 model of aggressive PDAC with prominent metastases to visceral organs^44,45^ revealed selective enrichment of iRFP720 signal in tumor-bearing mice (**Fig. 1a, b**) following a single intravenous (i.v.) dose of 2.5 x 10^7^ CFU. Tumor-selective CRC2631 colonization was confirmed by immunostaining demonstrating dense bacterial accumulation in the tumor core (**Fig. 1f, g**), but not in control pancreatic tissue (**Fig. 1d, e**). Quantitative chloramphenicol-based recovery showed over a 100-fold enrichment of CRC2631 in PDAC tumors and metastases relative to the liver or peripheral blood (**Fig. 1c**).

**Figure 1.**
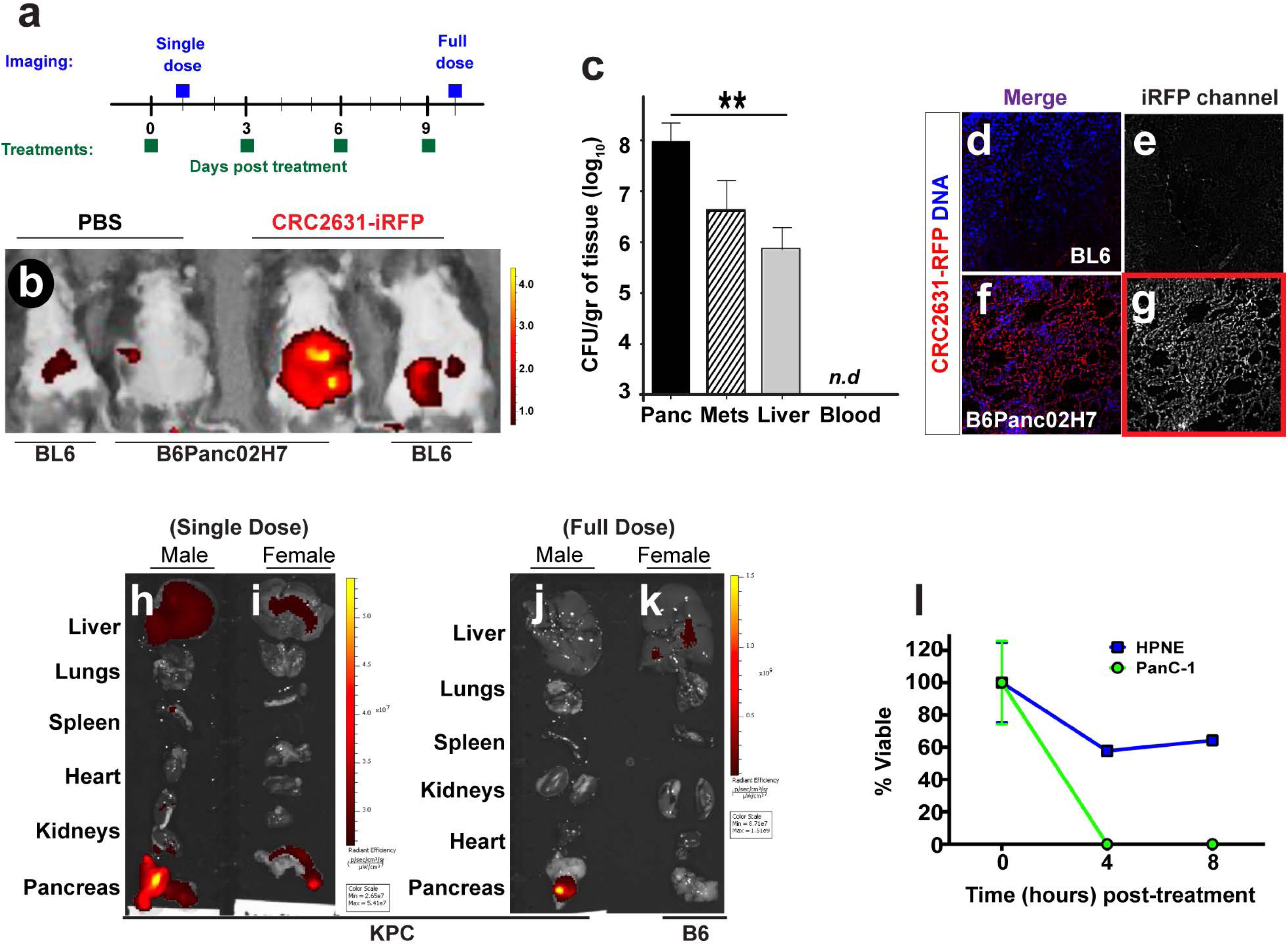
CRC2631 safely targets pancreatic tumors i*n vivo*. (**a**) Schematic indicating treatment and imaging schedules. (**b**) Representative *in vivo* fluorescence imaging images showing the biodistribution of iRFP-labelled CRC2631 (CRC2631-iRFP) in C57BL/6 (BL6) tumor-free controls or in orthotopic mouse PDAC model (B6Panc02H7) one day after intravenous administration of 2.5 x 10^7^ CFU and scanned. CRC2631-iRFP shows robust selective in B6Panc02H7 tumor-bearing mice compared to tumor-free controls. (**c**) Quantitative biodistribution confirms preferential accumulation of CRC2631 in primary tumors and visceral metastases ∼9 days after treatment. Collected tissues were weighed, homogenized, and dilutions were plated on chloramphenicol growth plates to selectively isolate CRC2631-iRFP counts per gram of tissue. CRC2631-iRFP counts/gr were ∼100-fold higher in tumor tissues than in the liver, a non-target clearing organs (**=*p<0.0077*). CRC2631-iRFP was undetectable in blood samples, suggesting low risk for CRC2631 translocation. (**d-g**) Representative fluorescence microscopy images of pancreases harvested from tumor-free C57BL/6J (BL6) controls or Panc02H7 mice ∼9 days after a single intravenous bolus of CRC2631-iRFP (*N=* 15/group). Pancreases were cross sectioned, counterstained with DAPI to detect nuclei, and examined under a fluorescent microscope. CRC2631-RFP signal (red) was enriched in tissues derived from in B6Panc02H7 orthotopic pancreatic tumor tissue compared to tissues from tumor-free controls. (**h-k**) **CRC2631-iRFP biodistribution in the genetic PDAC mouse model KPC** (Kras^G12D/+^; Trp53^R172H/+^; Pdx-1-Cre). C57BL/6J or MRI-verified tumor-positive KPC mice were treated intraperitoneally with a single or repeated (full) doses of CRC2631-iRFP (5 x 10^7^ CFU) as shown in (**a**). The indicated organs were harvested and imaged 1day post the first or the 4^th^ bolus corresponding to single or full dose schedule (**h, i or j, k**). CRC2631-iRFP selectively and persistently home to primary pancreatic tumors after both single or repeated (4x) dosing regimens. Tumor-free controls (**k**) show negligible signal. (**l**) Normal pre-cancerous (HPNE) versus cancerous pancreatic (PANC-1/PDAC) cells were treated with CRC2631-iRFP using a 4:1 bacterial to human cell ratio or multiplicity of infection/MOI. MTT assay shows CRC2631-iRFP preferentially kills PANC-1 cells.

We next evaluated CRC2631 tumor-targeting capabilities in the genetically engineered KPC model (Kras^G12D/+^; Trp53^R172H/+^; Pdx-1-Cre), which recapitulates hallmark features of human PDAC including fibrosis, immune exclusion, and therapeutic resistance^46–48^. Dose-escalation studies (5 x 10^7^ - 2 x 10^8^ CFU, intraperitoneal /i.p. or subcutaneous/s.c.) identified 5 x 10^7^ CFU i.p. every three days (x4 doses) as the best-tolerated regimen with limited weight loss, necropsy abnormalities, or treatment-related lethality (**Supplementary fig. 1**). As in the orthotopic model, CRC2631 persisted selectively within KPC tumors following single or repeated dosing, whereas the liver showed only transient early accumulation consistent with clearance (**Fig. 1h-k**). Similar tumor tropism was observed in mouse models of aggressive prostate cancer^35^ and immunotherapy-refractory Non-small Cell Lung Cancer (NSCLC) [Kras^G12D^ Tp53^+/-^L kb1^-/-^ (KPL)]^49^ (**Supplementary fig. 2a**). Consistent with tumor selectivity, CRC2631 preferentially killed PanC-1 PDAC cells compared to controls non-PDAC pancreatic cells (HPNE) (**Fig. 1l**).

Together, these results demonstrate that, in addition to prostate and lung cancer, CRC2631 safely and selectively colonize PDAC tumors.

### CRC2631 stimulates T-cell activation but induces a compensatory myeloid-recruiting program

To determine how CRC2631 alters tumor-intrinsic immune signaling in surviving cells, we performed RNA-seq on surviving Panc cells exposed to CRC2631 at a multiplicity of infection of 4 for up to 1hour. CRC2631 rapidly induced transcripts associated with lymphocyte trafficking and adhesion (CX3CL1 and ICAM1) alongside the myeloid chemoattractant CCL2 (**Fig. 2a**). These early transcriptional shifts suggest that CRC2631 simultaneously promotes effector T-cell recruitment while activating a counter-regulatory myeloid-recruiting program.

**Figure 2.**
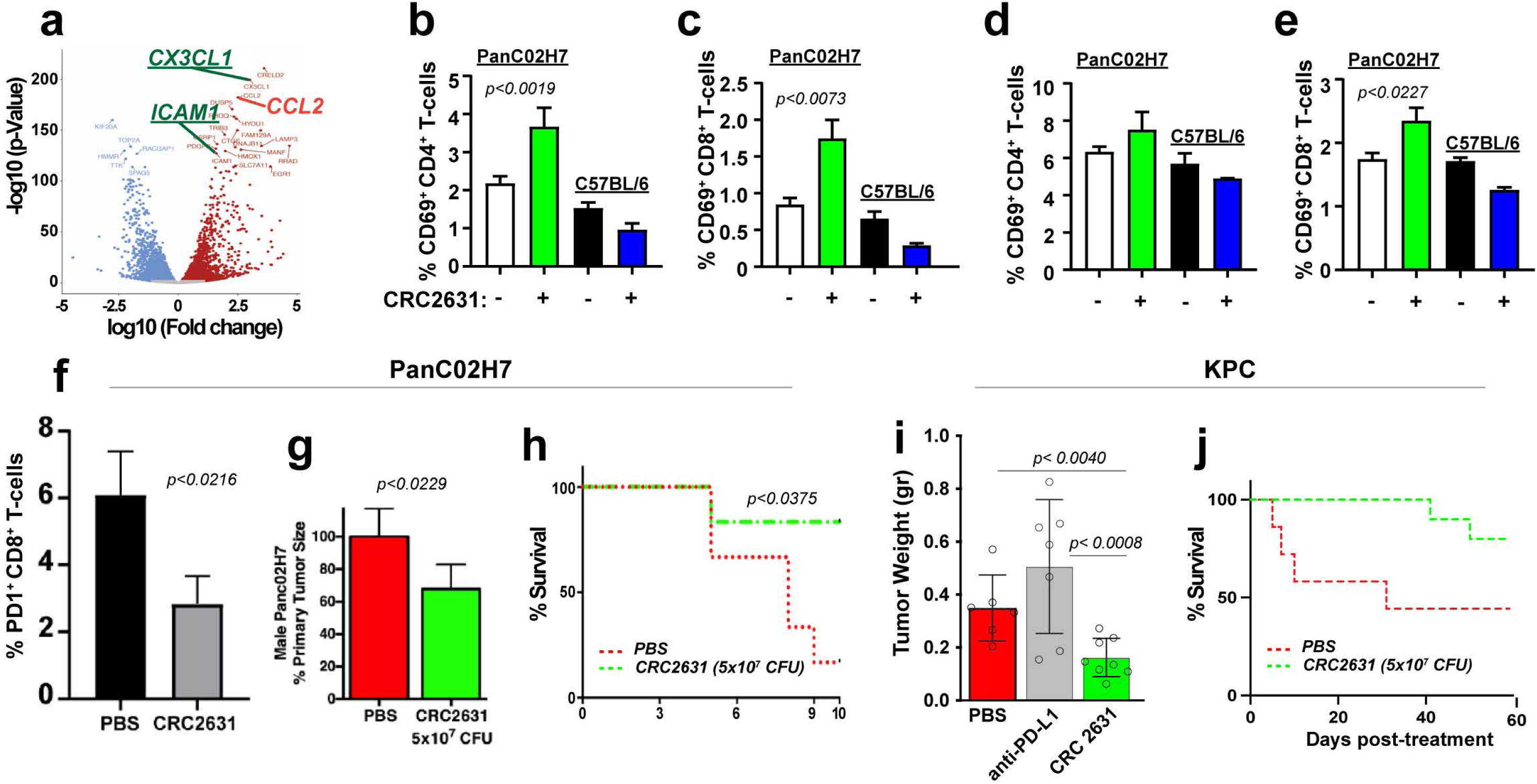
CRC2631 triggers tumor-specific immune responses and suppresses tumor growth. (**a**) Next-generation sequencing analysis of live Panc02 cells treated with PBS vs. CRC2631 (10^3^ MOI, 1 hour) shows upregulation of immune modulating molecules (CX3CL1, ICAM1, CCL2) compared to untreated controls. (**b-f**) flow cytometry immune profiling plots of size-controlled spleen from males (**b, c, *N= 7***) or female mice (**d, e, *N= 8***) mice or pancreatic tissues (**f**) harvested from C57BL/6J tumor-free controls or orthotopic PDAC mice (B6-Panc02H7). Tissues were harvested 8/9 days after intravenous treatment with a single bolus of CRC2631 (2.5 x 10^7^ CFU) or vehicle control (PBS). CRC2631 elevates the proportion of activated splenic (CD69+) CD4+ or CD8+ T-cells specifically in tumor-bearing mice. Also, CRC2631 decreased the abundance of exhaustion-prone (PD1-postive) CD8 T-cells (**f**). (**g, h**) CRC2631 reduces PanC tumor weight (*N=* 10, *p<*0.0229), and extended animal survival beyond the 10 days when most control mice die in the aggressive orthotopic Panc02H7 model (*N=* 10, *p<*0.0375). (**i, j**) Average tumor weight of pancreases harvested from KPC mice treated intraperitoneally with saline control (PBS) or anti-PD-L1(0.2 mg/mouse) or CRC2631 (5 x 10^7^ CFU). Anti-PD-L1 and CRC2631 were administered every 3 days, 4 doses total. CRC2631, but not anti-PD-L1, reduced tumor size (*N=* 8, *p<*0.0040) and extended survivorship (**j**). Thus, CRC2631 exhibit therapeutic effects in both, the syngeneic Panc02H7 and the KPC mouse models of pancreatic cancer. Error bars show standard dev. p-values are derived from student t-tests (**b-g**) or one-way ANOVA (**i**).

*In vivo*, CRC2631 increased splenic CD69^+^ CD4^+^ and CD69^+^ CD8^+^ T-cell populations in PDAC-bearing mice (**Fig. 2b-e**), but not in tumor-free animals, indicating tumor-dependent immune activation. Within PDAC tumors, CRC2631 increased total CD8⁺ T cells and reduced dysfunction-prone PD-1^+^ CD8^+^ subsets (**Fig. 2f**). These immune changes corresponded with ∼30% tumor reduction and improved survival in both the Panc02H7 orthotopic and KPC GEMM models (**Fig. 2g-j).**

Importantly, CRC2631 achieved tumor control under conditions where PD-(L)1 blockade failed to slow PDAC progression (**Fig. 2i**). Given PDAC’s profound resistance to checkpoint blockade, CRC2631’s ability to reduce tumor burden where PD-(L)1 inhibition is ineffective underscores its capacity to initiate immune activation through mechanisms not accessible to checkpoint therapy alone.

Despite these beneficial shifts, intratumoral CD8^+^ T cells remained PD-1^+^ and activated (CD69⁺) CD8⁺ cells diminished overtime, suggesting incomplete reversal of CD8^+^ exhaustion.

Pathogen- and damage-associated molecular patterns (PAMPs/DAMPs) from intratumoral bacteria are well known to engage pattern-recognition receptors such as Toll-like Receptors (TLRs) on tumor and myeloid cells, activating canonical NF-κB and type I/II interferon pathways and driving production of chemokines including CCL2 that recruit inflammatory monocytes/MDSCs to sites of infection or tissue stress^50–53^. In line with this paradigm, bulk RNA-seq of Panc02 cells exposed to CRC2631 shows rapid induction of *Ccl2* together with the lymphocyte-trafficking and adhesion genes (**Fig. 2a**). Further, spatial transcriptomic profiling of KPL NSCLC tumors treated with CRC2631 not only revealed transcriptional shifts consistent with lymphocytes trafficking and adhesion, but also robust activation of a TLR4-NF-κB cascade, type I/II IFN signaling, and antigen-processing/MHC-I machinery (*Tap1, Psmb8/9, H2-K1/H2-D1, B2m*), together with increased *Cd274* (PD-L1) (**Supplementary fig. 3a-t**). These immune responses were accompanied with an upregulation of cell death transcriptional program in colonized tumors (**Supplementary fig. 2a, b and supplementary fig. 3u**). However, immune cell-types deconvolution of the spatial transcriptomic data showed enrichment of Ly6C^high^ MDSCs signatures following CRC2631 treatment in KPL mice (**Supplementary fig.4e**), mirroring emergence of immunosuppression observed in PDAC mice.

Taken together, these observations are consistent with CRC2631 functioning as a potent intratumoral danger signal that enhances antigen presentation yet reinforces a myeloid bottleneck.

We sought to overcome this myeloid barrier. The CKLF-like MARVEL transmembrane domain containing 6 (CMTM6) protein has emerged as a central regulator of tumor immune evasion^14–23^: genome-wide CRISPR screens identified CMTM6 as a critical stabilizer of PD-L1 on tumor and myeloid cells, such that CMTM6 loss reduces surface PD-L1 and restores tumor-specific T-cell activity^15,21^. More recent studies demonstrate that genetic suppression of tumor- or host-intrinsic CMTM6 drives potent CD8⁺/NK-dependent antitumor cytotoxicity even when the PD-1/PD-L1 axis is genetically ablated, establishing a PD-L1-independent checkpoint function^17^. Further, tumor-intrinsic CMTM6 drives M2-like macrophage polarization and the formation of immunosuppressive myeloid niches^22,23^. High CMTM6 expression in the TME correlates with myeloid-rich immune cold tumors and poor prognosis in PDAC and other solid cancers^18–20^.

Therefore, we engineered iSTORM, a CRC2631 derivative expressing *sh-Cmtm6*, to preserve CRC2631-mediated PAMP/DAMP-NF-κB/IFN activation yet dismantle CMTM6-medialted lymphoid and myeloid checkpoints, converting the bacteria-induced myeloid response into a more robust CD8⁺/NK-dominant antitumor state.

### iSTORM safely enhances T-cell/NK activation and suppresses myeloid recruitment

We first assessed iSTORM safety and target engagement. Comprehensive acute and chronic toxicity studies, including blood chemistry, immune profiling, and histopathology in C57BL/6 or FVBnj mice confirmed that iSTORM maintains its favorable safety profile (**Fig. 3 and Supplementary fig. 5**).

**Figure 3.**
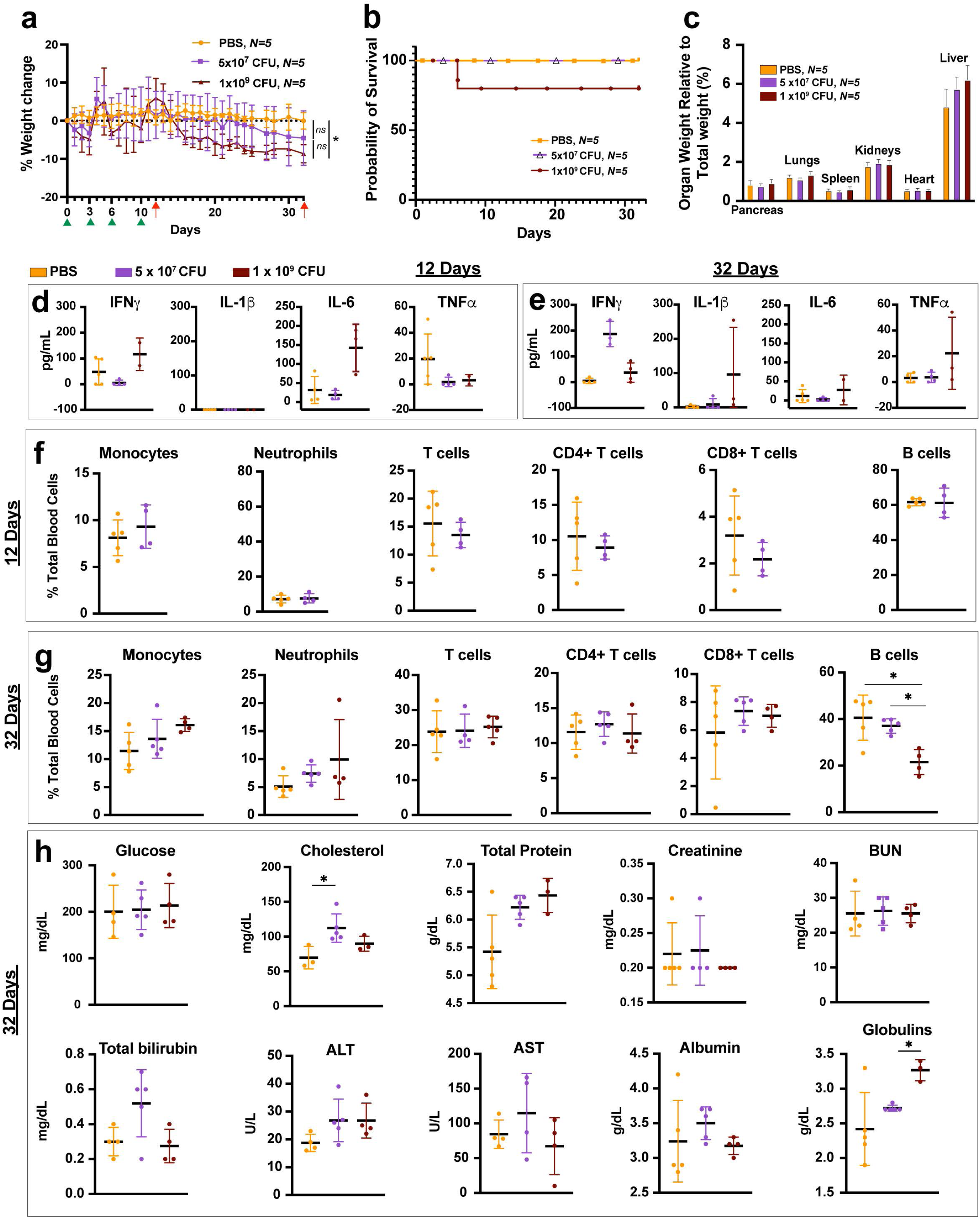
Safety and tolerability of iSTORM in C57BL/6 mice. 13-15-weeks old female C57BL/6J mice received four intravenous doses of saline control (PBS) (yellow, *N= 5*) or iSTORM on days 0, 3, 6, and 10 (green arrowheads) at either 5 x 10^7^ CFU (purple, *N= 5*) or 10^9^ CFU (brown, *N= 5*). Following treatment, Body weight was recorded daily and expressed as percent change relative to baseline (day 0) (**a**), which allowed to observe a progressive weight loss in mice from 10^9^ CFU arm in respect to the vehicle, but still inferior to 10% across the whole duration of the study. Survival (**b**) was monitored for 32 days, at which point indicated organs were harvested, weighed, and normalized to total body weight (**c**). Error bars show standard deviation and p-values are derived from two-way repeated measure ANOVA (**a**), Log-Rank test (**b**) and Student’s t-test (**c**). (**d, e**) Blood from the animal cohort above was collected on day 12 or 32 (red arrows) to assess the acute and longer-term impact of CRC2631 on plasma cytokines levels. Error bars show standard deviation and p-values are derived from Student’s t-test. (**f-h**) Serial immune cell counts (**f, g**) or organ function chemistry (**h**) were assessed on day 32, and on days 12 and 32, respectively. Blood chemistry showed a statistically significant increase (*p<*0.05) in (1) hematic globulins between 10^9^ CFU arm and 5 x 10^7^ CFU arm, and (2) cholesterol between 5 x 10^7^ CFU arm and the vehicle (*p<*0.05), and (3) a decrease in B cells across all the treatments in respect to the vehicle. Error bars show standard deviation and p-values are derived from Student’s t-test.

As expected, iSTORM reduced CMTM6 and PD-L1 in PDAC cells (**Fig. 4a, b**), maintained its tumor-targeting capabilities *in vivo* (**Fig.4c-g**), and significantly increased activated (CD69^+^) CD8^+^ T cells within PDAC tumors compared to CRC2631 or PBS in the genetic KPC mouse model of PDAC (**Fig. 4h**). Consistent with enhanced immune re-potentiation in PDAC, spatial transcriptomic analysis in the KPL mouse model of immunotherapy resistant NSCLC revealed that iSTORM reduces CMTM6 mRNA (**Supplementary fig. 4a**). Although both CRC2631 and iSTORM trigger interferon-γ and granzymes transcriptional signatures compared to saline controls (**Supplementary fig. 4 h-k)**, iSTORM shows more robust immune responses: it elevates antigen processing/display and infiltration of NK and NKT while reducing suppressive myeloid subsets in the TME more robustly than CRC2631 **(Supplementary fig. 4b-g).** Consistent with therapeutic immune remodeling in the NSCLC KPL model, iSTORM significantly reduced tumor growth relative to PBS control **(Supplementary fig. 4l).** Taken together, these data show that iSTORM deepens myeloid suppression and enhances antitumor immunity.

**Figure 4.**
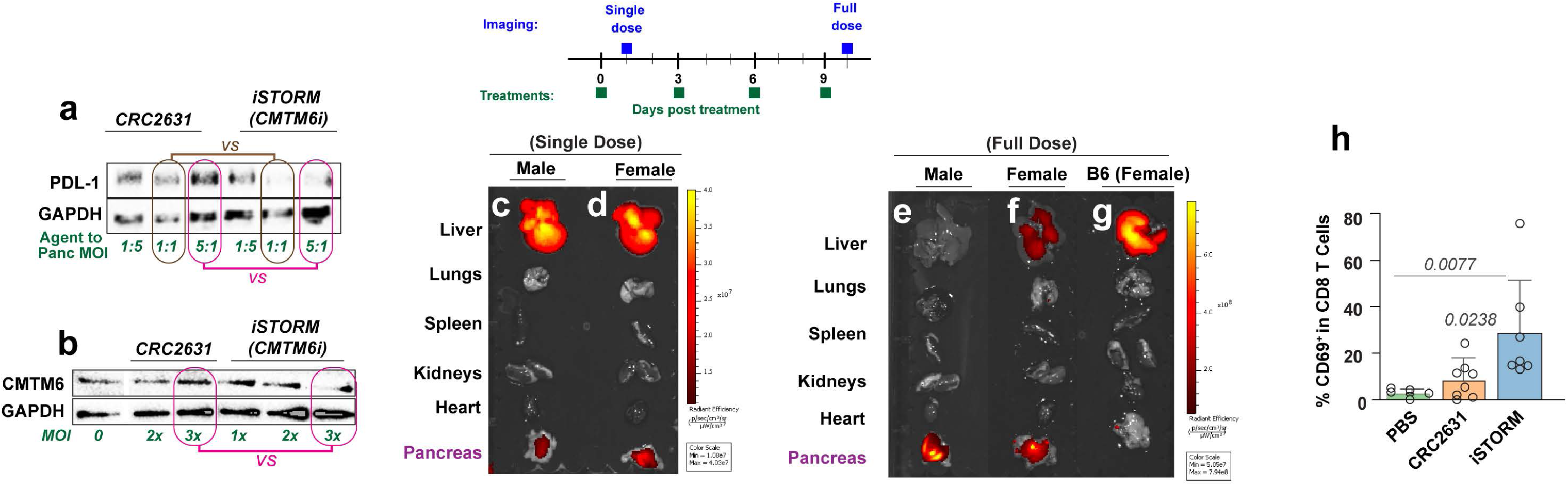
iSTORM depletes PD-L1/CMTM6 and enhances CD8 activation in pancreatic tumors. (**a, b**) iSTORM reduces PDL-1 and CMTM6 protein levels in PDAC cells. Intact Panc02 cells or Panc02 cells overexpressing PDL-1 were treated with CRC2631 (lanes 1-3) or iSTORM (CRC2631 expressing shRNA against CMTM6, lanes 4-6) at the indicated bacteria/PanC cell ratios. Living cells were isolated and lysed for Western blot. Blots were stained against CMTM6 or PDL-1 or GAPDH. (**c-g**) Representative Images from ex-vivo IVIS fluorescence imaging of indicated organs harvested from MRI-verified tumor-bearing KPC GEMM mice treated with a single (**c, d**) or 4 doses (**e, f**) of iSTORM (5 x 10^7^ CFU), each administered every 3 days. Additional tumor-free C57BL/6J (B6) specificity controls were included (**g**). iSTORM transiently accumulates in the liver but persists in tumor tissues after full dose regimen, specifically in tumor-bearing mice. (**h**) Representative graph showing comparative abundance of CD69^+^/CD8^+^ T-cells by flow cytometry (FACS). Pancreases were harvested from PDAC bearing KPC mice treated intraperitoneally with a complete dose of saline control (PBS) or 5 x 10^7^ CFU of either CRC2631 or iSTORM (4 doses, 1 every 3 days). iSTORM increases the abundance of CD69^+^/CD8^+^ T-cells much more potently than CRC2631.

To standardize dosing regimen and support translational scalability, we generated a lyophilized formulation (iSTORM-L) that maintains therapeutic efficacy without cold chain production. Toxicity studies in mice confirmed that iSTORM-L (5 x 10^7^ or 5 x 10^8^ CFU) maintains a favorable safety profile based on weight and immune profiling (**Supplementary fig. 6a, b**).

We assessed iSTORM-L capability to generate immune and tumor control efficacy signals. Comparative growth kinetic analyses of the lyophilized strain versus the cultured counterpart revealed that reconstituted lyophilized bacteria expand more slowly than cultured stocks over the first 10 h, reaching only ∼70-80% of the optical density (OD) achieved by cultured bacteria at the same starting CFU (**Supplementary fig. 7**). Because early intratumoral expansion is likely to be a major determinant of therapeutic activity, we reasoned that a nominally identical inoculum would under-represent the lyophilized product. We therefore evaluated two lyophilized doses *in vivo*: (i) 5 x 10^7^ CFU, matching the dose used for cultured bacteria, and (ii) an increased dose of 1 x 10^8^ CFU to compensate for the early growth deficit and safely detect dose-dependent effects.

Peripheral blood immune profiling revealed that iSTORM-L fully preserves the immune-stimulatory activity of the original strain, producing a clear, graded immune activation program across dose levels. Escalating iSTORM-L doses expanded circulating CD44^+^CD69^+^ and ICOS^+^CD4^+^ T cells while reducing PD-1^+^ and PD-1^+^LAG3^+^ exhaustion-prone subsets (**Fig. 5a**). CD8^+^ T cells showed a parallel activation pattern with a strong expansion of ICOS⁺ effector-like CD8^+^ T cells and uniformly low PD-1^+^ or PD-1^+^TIM3^+^ populations across doses (**Fig. 5b**).

**Figure 5.**
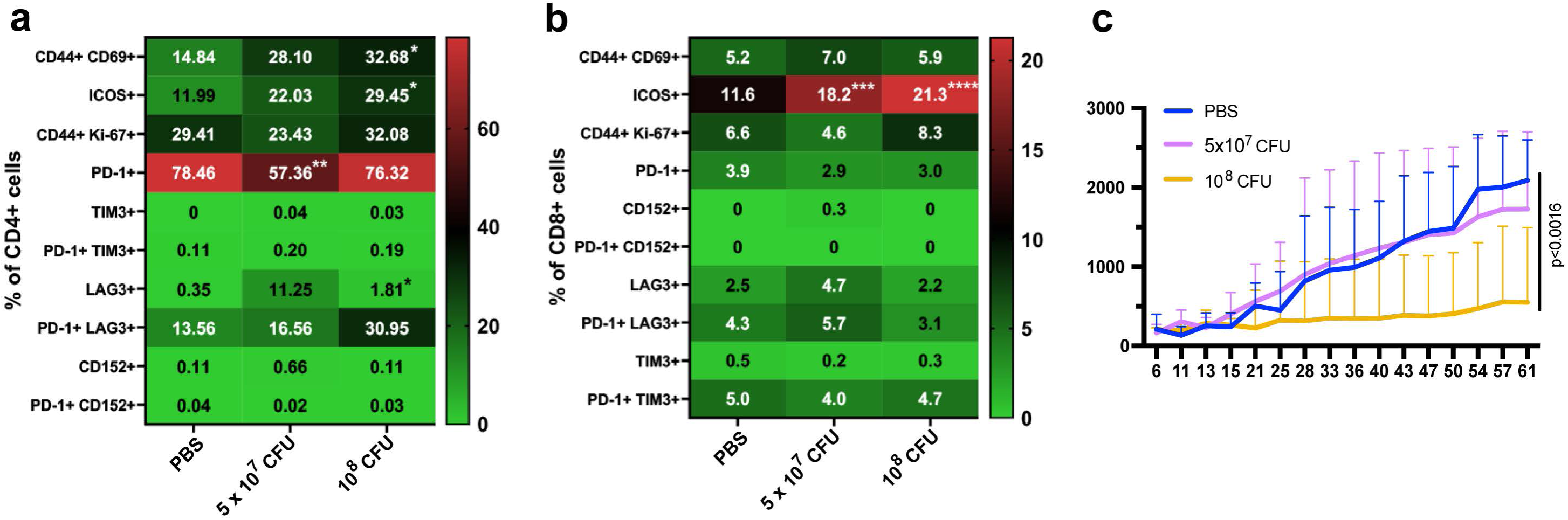
Lyophilized iSTORM activates immune cells and achieves tumor control. (**a, b**) Peripheral blood immune profiling in the KPC flank model. KPC cells (1.0 x 10^6^) cells were inoculated into the right flanks of 6-8-week-old C57BL/6J mice. Once tumors reach the target volume of 80-120 mm³ mice were randomized to treatment group (n=8/group) and treated with PBS or lyophilized iSTORM (iSTORM-L) at 5×10^7^ CFU or 1×10^8^ CFU/mouse, administered intraperitoneally, one bolus every 3 days, 4 doses total). Immune profiling in peripheral blood collected 15/16 days post the last treatment (23/24 days after the first dose) demonstrates that iSTROM fully retains immune-stimulatory activities and elicits a graded, dose-dependent activation of circulating T-cell compartments. Increasing iSTORM-L doses expanded activated CD4^+^ T cells (CD44^+^CD69^+^ and ICOS^+^) while concomitantly reducing exhaustion-associated PD-1^+^ and PD-1^+^LAG3^+^ CD4^+^ subsets (**a**). CD8^+^ T cells displayed a parallel activation program, characterized by robust expansion of ICOS⁺ effector-like CD8^+^ T cells and uniformly low frequencies of PD-1^+^ or PD-1^+^TIM3^+^ exhausted populations across all doses (**b**). **c**) iSTORM-L induced dose-dependent tumor growth suppression in the KPC mice above. Tumor volumes (caliper measurements) were measured twice per week. Bar and whisker represent means and standard deviation, respectively.

Consistent with its immunostimulatory effect, iSTORM-L produced a robust dose-dependent tumor suppression in the KPC flank model. The 5 x 10^7^ CFU iSTORM-L dose failed to control tumor growth, despite enhanced immune signatures, whereas the highest dose (1 x 10^8^ CFU) suppressed tumor growth (**Fig. 5c**). The highest dose reduced tumor expansion by more than 50%, showing significant benefit over both control and lower doses. These findings confirm that the lyophilized formulation maintains full biological potency.

## DISCUSSION

Pancreatic ductal adenocarcinoma (PDAC) remains a paradigm of treatment resistance, where the failure of anti-PD-1 therapy underscores a fundamental challenge: the dense stroma limits drug and immune penetration, and myeloid-dominated suppression extinguishes cytotoxic activity, leading to the failure of most systemic immunotherapies^7–9,12,13^. This closed and hostile tumor ecosystem has rendered most systemic immunotherapies ineffective. Here, we show that the tumor-colonizing microbial platform CRC2631, and its engineered derivative iSTORM, directly dismantle these barriers by achieving selective intratumoral colonization, initiating upstream T-cell activation, and disrupting the CMTM6-dependent circuitry that stabilizes PD-L1 and orchestrates myeloid immune suppression.

CRC2631 is fundamentally distinct from previous bacterial cancer therapies. Its lineage, shaped by decades of nutrient-restriction evolution and refined through rfaH deletion and aroA/thyA auxotrophy, confers intrinsic tumor selectivity and a markedly improved safety profile over first-generation agents like VNP20009^35–40^. Consistent with these properties, CRC2631 exhibited >100-fold enrichment in orthotopic and spontaneous PDAC tumors with minimal off-target persistence and toxicities.

Direct *in vitro* killing assays confirmed that CRC2631 selectively eliminates human pancreatic cancer cells, while sparing non-malignant controls at matched MOI (**Fig.1l**). We propose that this selectivity arises not from engagement of a single canonical receptor, but primarily from recognition of a tumor-restricted surface glycoprotein and glycan state. Indeed, high-throughput glycan binding profiling demonstrated that CRC2631 preferentially recognizes mannose-rich and hybrid N-glycans, including oligomannose structures associated with transferrin and transferrin receptor (TfR)-linked glycoforms (e.g., Man_5_-Man_9_)^54^, exhibiting up to ∼400-fold higher binding signal compared with non-mannosylated glycans (**Supplementary fig. 8a, b**). This binding data indicates a strong preference for incompletely matured N-glycan architectures, a feature widely reported to accompany oncogenic stress, hypoxia, and immune exclusion in solid tumors^55,56^.

Consistent with this targeting mechanism, TfR was overexpressed in PDAC cell lines (PANC-1, CFPAC-1) relative to non-malignant HPNE controls and exhibited a PNGase-sensitive mobility shift, confirming N-glycosylation (**Supplementary fig. 8c, d**). Functional blockade of TfR using antibodies significantly reduced iSTORM-mediated cytotoxicity in PANC-1 and KPC cells (**Supplementary fig. 8e**). This partial dependence is consistent with a multivalent targeting mechanism, in which CRC2631/iSTORM engages a shared glycosylation program displayed across multiple tumor-associated glycoproteins rather than relying on a single receptor interaction or mechanism.

This glycosylation state, surface-exposed high-mannose and incompletely processed N-glycans^55,57^, is selectively enriched in immune-cold, fibrotic malignancies such as PDAC ^57–61^ and non-small cell lung cancers ^62,63^, which are refractory to immune checkpoint blockade^62,64^. In this context, CRC2631 activated systemic and intratumoral T-cell responses and mediated antitumor effects in settings where anti-PD-1 therapy was ineffective, supporting a model in which iSTORM overcomes immune exclusion by combining upstream immune activation with CMTM6-dependent disruption of tumor and myeloid immune-suppressive programs.

Together, these findings support a model in which iSTORM exploits tumor-specific nutrient receptor glycosylation and glycan convergence states to anchor within immune-excluded tumor cores, penetrate fibrotic tissue, and initiate immune activation independently of canonical checkpoint sensitivity. Future *in vivo* genetic and glycan-editing studies targeting mannose-rich N-glycan will be essential to quantitatively define the contribution of this targeting mechanism to intratumoral colonization and therapeutic efficacy.

CRC2631 revealed a compensatory response intrinsic to PDAC. A surge in myeloid recruitment, possibly driven by CCL2 induction (**Fig. 2a** ) and reflected in expanded MDSC signatures (**Supplementary fig. 4e and** ^65–68^). This bottleneck may explain why bacterial therapies that merely induce inflammation yield limited durability. iSTORM is engineered to overcome this limitation by delivering the first tumor-localized CMTM6 blockade, reducing PD-L1 stability and disrupts CMTM6-mediated myeloid recruitment^14–20^, deepening CD8⁺ T-cell activation and depleting bone-marrow-derived MDSCs. Future studies will elucidate immune memory, TCR clonality, and antigen spread in metastatic settings.

This precise immune rewiring provides a rationale for combination therapy approaches. Because iSTORM inhibits CMTM6/PD-L1 yet a residual MDSCs subpopulation persists, agents that selectively target these subsets (e.g., gemcitabine, CSF1R inhibitors, or anti-Ly6G antibodies) can be leveraged to remove this potential bottleneck and maximize anti-tumor immune activities. Also, direct engineering of CCL2 inhibition into iSTORM/CRC2631 may deepen and prolong anti-tumor immune activities.

iSTORM therapeutic potential benefits from a multi-layered safety architecture. LPS attenuation, auxotrophy that ensure replication preferentially in tumors, rapid and safe hepatic clearance, and defined sensitivity to established antibiotics, providing a reliable antibiotic kill-switch. Also, the lyophilized formulation (iSTORM-L) ensures clinical practicality, conferring long-term stability, cold-chain independence, and consistent, dose-dependent immune activation, all vital for global deployment.

In summary, this study introduces a microbial immunotherapy platform that integrates multi-mechanistic tumor targeting, first-in-class CMTM6 blockade, and clinical readiness through lyophilization and built-in biocontainment. iSTORM’s ability to function where checkpoint inhibitors fail, to dismantle both PD-L1 stability and myeloid suppression, and to do so with a scalable, cost-effective format, positions it as a transformative candidate for PDAC and other solid immunologically cold tumors.

## ETHICS APPROVAL

All animal experiments were conducted in accordance with institutional guidelines and were approved by the University of Missouri Institutional Animal Care and Use Committee (IACUC MU protocol #63841). Mice were housed in an AAALAC-accredited facility, and all procedures adhered to established humane endpoints. This study used only established human cell lines obtained from commercial or institutional repositories; no human subjects, human tissue, or identifiable human data were involved.

## CONSENT FOR PUBLICATION

All authors have read and approved the final version of this manuscript and consent to its publication.

## AVAILABILITY OF DATA AND MATERIAL

All data supporting the findings of this study are included in this article and its supplementary information. Raw datasets and materials, including RNA-seq count matrices, flow cytometry files, and glycan array readouts, are available from the corresponding author on reasonable request. Plasmids and bacterial strains may be made available under a material transfer agreement with the University of Missouri.

## AUTHORS CONTRIBUTIONS

**Gunter K:** coordinated mouse studies and colony maintenance, including tail biopsies for genotyping, therapeutic treatments, tumor harvesting and weighing, survivorship monitoring, and animal weighing.

**Kazmierczak R:** constructed and validated bacterial strains and performed biodistribution studies in the orthotopic Panc01H7 PDAC model.

**Hasani M:** carried out animal maintenance, tail biopsies for genotyping, tamoxifen induction, therapeutic injections, coordination of MRI and IVIS imaging, tumor harvesting and weighing, and survivorship monitoring with regular body-weight measurements.

**Patterson N:** coordinated overall mouse work, generated the genetic KPC mouse model, supervised therapeutic treatments and colony maintenance, and harvested and processed tumor samples for flow cytometry and ex vivo imaging.

**Wang Q:** together with **Li G**, generated the orthotopic Panc01H7 PDAC model and performed initial analyses of treatment-induced immune shifts.

**Canti L**: performed iSTORM toxicity studies, including clinical monitoring of mice, euthanasia, submission of plasma and tissue samples for external histopathologic scoring, and bacterial growth-kinetics assays.

**Tesfay M.Z:** assessed tumor size responses and therapeutic efficacy of iSTORM-L.

**Cios A**: performed peripheral blood immune profiling in iSTORM-L–treated cohorts.

**Dhagat B**: performed next-generation sequencing profiling of PDAC cells.

**Pastor M.Q**: carried out iSTORM biodistribution and tumor control efficacy studies in KPL mice.

**De la Nuez C**: performed transferrin receptor (TfR) expression and glycosylation analyses and TfR functional studies.

**Verburg T**: validated iSTORM-mediated CMTM6 inhibition in PDAC cells.

**Moyer J**: performed animal maintenance, tail biopsies for genotyping, therapeutic treatments, tumor induction and harvesting, tumor weight measurements, and survivorship monitoring with regular body-weight measurements.

**Mwanza M**: carried out animal maintenance, therapeutic treatments, macroscopic necropsy-based toxicity assessments, and organ harvesting.

**Moaven O**: contributed to study conceptualization, formal data analysis, and manuscript review and editing.

**Li G**: generated the orthotopic Panc01H7 PDAC model and performed initial analyses of treatment-induced immune shifts.

**de Figueiredo P**: contributed to study conceptualization, data analysis, and manuscript review and editing.

**Nagalo B.M**: contributed to study conceptualization, data curation and formal analysis, and manuscript review and editing.

**Chabu CY**: led study conceptualization, data curation and analysis, funding acquisition, project administration, original manuscript drafting, and manuscript review and editing.

## ACKNOWLEDGEMENT

We thank the Office of Animal Resources (OAR), University of Missouri, for technical support with animal studies and Megan Polniak (University of Missouri) for assistance with colony maintenance. We are grateful to Steven Dubinett (UCLA) for providing KPL cells, OPS Diagnostics for performing lyophilization, and Bruker Spatial Biology for support with spatial transcriptomics. We also thank the Emory University Consortium for Functional Glycomics (Atlanta, GA) for lectin array resources.

## MATERIAL AND METHODS

### Bacteria work

Isolated colonies of bacteria were grown from -80°C stock aliquots frozen in 25% glycerol (Fisher, catalog# BP229-1, MA, United States) on solid or liquid LB media (Fisher/Research Products International, catalog# M342-500, MA, United States) supplemented with 200 μg/mL thymine (Arcos Pharmaceuticals) and antibiotics: 50 μg/mL kanamycin (Sigma, catalog# 60615, MO, United States), 50 μg/mL ampicillin (Sigma, catalog# A9393) or 20 μg/mL chloramphenicol (Gold Biotechnology, catalog# C-105-5, MO, United States), as required by the strain. Bacteria grown on solid media was incubated for 24-30 h at 37°C before use. Liquid media cultures were incubated in 50 mL sterile tubes for 20-24 h in a 37°C, 220 rpm dry shaking incubator. Strains grown for injection were washed with sterile phosphate buffered saline (PBS) (Rocky Mountain Biologicals, catalog# PBS-BBZ, MT, United States) and concentration adjusted for injection and for *in vitro* cell viability assays.

#### Engineering of iSTORM-v1 (CRC2631-iRFP720 carrying pLKO.1 shRNA against CMTM6)

CRC2631-iRFP720 was described previously^35^. A short hairpin RNA targeting CMTM6 (shCMTM6) was cloned into the pLKO.1 backbone (Addgene, Watertown, MA, plasmid #10878). The resulting plasmid was introduced into CRC2631-iRFP720 by electroporation. Briefly, electrocompetent cells were prepared by repeated washing in ice-cold sterile 10% glycerol and mixed with 50-100 ng of plasmid DNA. Electroporation was performed in 0.2 cm gap cuvettes at 2.5 kV, 25 µF, and 200 Ω. Immediately following pulsing, cells were recovered in pre-warmed LB medium and incubated at 37°C with shaking for 1-2 h. Bacteria were plated on LB agar supplemented with thymine (200 µg/mL) and ampicillin 100 µg/mL for selection. Positive transformants were screened, and correct plasmid incorporation and shCMTM6 cassette sequence integrity were confirmed by Sanger sequencing prior to downstream experiments.

#### Preparation of cultured iSTORM/CRC2631-iRFP

Cultured iSTORM was grown by picking a single colony from an LB agar plate and letting it expand overnight in liquid LB. Liquid and solid bacterial media were prepared as described in the section above, including the appropriate antibiotic for selection. After O/N incubation, the OD_600_ of the culture was measured using a Nanodrop™ 2000c cuvette spectrophotometer (Thermo Fisher Scientific) and converted to CFU/mL based on standard curve as described previously^35^.

#### Growth kinetics of lyophilized vs cultured iSTORM

The contents of (1) one vial of lyophilized iSTORM reconstituted in PBS and (2) the surface layer of a 25% glycerol stock of cultured iSTORM were transferred into liquid LB containing the selection antibiotic and incubated for 8 hours (shaking at 220 rpm, at 37°C) to maintain bacteria in log phase. The cultures were then transferred at a density corresponding to 0.1 OD_600_ in a 96-well plate and incubated at 37°C with constant orbital shaking in a Synergy H1 microplate reader (Agilent, CA, United States). Absorbance of bacterial cultures was automatically measured every 20 mins in a plate reader.

#### Glycan-dependent targeting assays

Lectin/glycan array profiling was carried out by the Consortium for Functional Glycomics at Emory University Atlanta GA. Briefly, bacterial binding specificity was assessed using CFG (Consortium for Functional Glycomics) glass glycan arrays (version 2.1) containing chemically defined glycans printed on N-hydroxysuccinimide-activated glass slides. Bacteria were grown overnight in LB supplemented with thymine (200 µg/mL) at 37°C with shaking to an OD600 of approximately 1.0 (approximately 1 x 10^8^ bacteria/mL), harvested by centrifugation, and washed twice with ice-cold PBS. Bacterial pellets were resuspended in PBS and stained with SYTO 83 (50 µM final concentration; Thermo Fisher Scientific, Waltham, MA, USA) for 1 h at room temperature with gentle mixing, then washed twice with cold PBS to remove unbound dye.

CFG array slides were pre-hydrated in Milli-Q water for 5 min and dried by centrifugation for 1 min. Labeled bacteria were overlaid onto slides and incubated for 1 h at room temperature. Slides were washed three times in cold PBS (30 seconds per wash, fresh buffer each wash), briefly rinsed in Milli-Q water, and dried by centrifugation for 1 min. Arrays were scanned using a 543 nm laser to quantify relative glycan recognition.

Fluorescence intensity was quantified as relative fluorescence units (RFU) for each spot. For each glycan, mean RFU, standard deviation, standard error of the mean (SEM), and percent coefficient of variation (%CV) were calculated. To reduce sporadic outlier-driven false positives, the highest and lowest RFU values within each set of six replicate spots were excluded prior to averaging; reported mean RFU values reflect the average of the remaining four replicates.

#### Functional validation of N-glycan requirement

Human pancreatic ductal epithelial control cells (HPNE) and pancreatic cancer cells (CFAC-1/ ATCC Catalog# CRL-1918, PANC1/ATCC catalog#CRL-1469, KPC/Kerafast catalog# EUP012-FP) were grown to 80-90% confluency in 6-well plates. Cells were co-cultured with CRC2631 or iSTORM-v1 at MOIs spanning 1:1 to 1:30 for 4-8 h at 37°C and 5% CO2. Extracellular bacteria were eliminated by adding gentamicin to a final concentration of 40 µg/mL (Fresenius Kabi, IL, USA; catalog# 401897G) for 1 h.

For PNGase assays, cells were lysed in Pierce RIPA buffer (Thermo Fisher Scientific; catalog# 89900). Lysates were treated with PNGase (Gibco, Thermo Fisher Scientific; catalog# A39245V) according to the manufacturer’s instructions. Transferrin receptor (TfR/CD71) abundance and PNGase-sensitive mobility shifts were assessed by immunoblotting with anti-TfR/CD71 (Developmental Studies Hybridoma Bank, Iowa City, IA, USA; catalog# G1/221/12; 1:500 dilution) and HRP-conjugated anti-GAPDH (Cell Signaling Technology, Danvers, MA, USA; catalog# 3683S; 1:1000 dilution). Membranes were developed using SuperSignal West Pico PLUS (Thermo Fisher Scientific; 34580) and SuperSignal West Femto (Thermo Fisher Scientific; catalog# 34095) mixed at a 1:50 ratio and imaged on a Gel Doc XR+ system (Bio-Rad, Hercules, CA, USA). Host cell entry-dependent cytotoxicity was assessed using MTT assays as previously described^35^.

#### Cell Lines, Medium, and Culture

Murine Panc02-H7 pancreatic cancer cells are an invasive cell line derived from Panc02^44^. Panc02-H7 cell lines were maintained in Dulbecco’s Modified Eagle Medium (DMEM) with 2 mmol/L L-glutamine, 10 mmol/L HEPES (Thermo Fisher Scientific), supplemented with 100 U/mL penicillin (Thermo Fisher Scientific), 100 mg/mL streptomycin (Thermo Fisher Scientific), and 10% FBS (Fetal Bovine Serum - Sigma-Aldrich, MO, United States). KPC cells (Kerafast, catalog# EUP012-FP, MA, United States) were maintained in DMEM-High Glucose (Gibco^TM^, catalog# 11965092, NY, United States) supplemented with 10% FBS (Sigma-Aldrich, catalog# F4135, MO, United States). KPL cells were maintained in RPMI (Gibco, catalog# 11875093), supplemented with 10% FBS (Sigma-Aldrich, catalog# F4135). All cell lines are mycoplasma-negative and grown at 37°C with 5% CO_2_ in a humidified atmosphere.

### Co-culture Assay

For co-culture assays, pancreatic cells were grown and harvested at 80-90% confluency. Cells were plated on 6-well plates and infected with iSTORM/CRC2631, grown in liquid LB media for 20-24 hours at 37°C and 220 rpm, using an MOI ranging from 1 to 30. ISTORM co-culture was incubated at 37°C, 5% CO_2_ for 4 hours, then gentamicin (Fresenius Kabi, catalog# 401897G, IL, United States) was added at a concentration of 40 μg/mL and left to incubate for an additional hour to eliminate extracellular bacteria. Cells were processed either for MTT viability assays or Western blotting.

#### Western blotting

Cells were lysed with Pierce^®^ RIPA buffer (Thermo Fisher Scientific, catalog# 89900). Transferrin receptor antibody, anti-CD71 (DSHB, catalog# G1/221/12, IA, United States) and -CMTM6 (Sigma, catalog# SAB2701009) were used at 1:500 and 1:5000 dilution, respectively, and GAPDH HRP-conjugated (Cell Signaling, catalog# 3683S, 1:1000 dilution) on PVDF membrane (Sigma-Aldrich, catalog# IPVH00010). In PNGase assay, lysates were treated with PNGase (Gibco^TM^, catalog# A39245V) as recommended by the manufacturer. Blots were developed using SuperSignal^TM^ West Pico PLUS (Thermo Fisher Scientific, catalog #34580) and SuperSignal^TM^ West Femto (Thermo Fisher Scientific, catalog# 34095) in a 1:50 ratio on a Gel Doc^TM^ XR+ Imager (Bio-Rad, CA, United States).

### Mouse work

All animal studies were conducted under protocols approved by the University of Missouri Institutional Animal Care and Use Committee (IACUC) and performed in accordance with institutional biosafety policies. Mice were housed under standard conditions (12 h light/12 h dark cycle) in individually ventilated cages with ad libitum access to water and standard diet 20 unless otherwise specified for imaging studies. Animals were euthanized upon reaching predefined humane endpoints, including tumor ulceration or tumor size exceeding 15 mm.

#### The B6Panc02-H7 pancreatic orthotopic model^45^

Briefly, 12-15 weeks old C57BL/6 mice were anesthetized and a left flank incision of 1.5 cm was made to open the peritoneal cavity. The spleen was manipulated to visualize the pancreas and 2.5 x 10^5^ Panc02-H7 cells in 20 µL PBS was injected into the pancreatic tail. All organs were returned to the peritoneal cavity, the incision sutured and skin clipped. Tumors and spleens were harvested, weighed, and processed for immune profiling by flow cytometry.

#### KPL-FVB/NJ subcutaneous tumor models in FVB/NJ immunocompetent mice

FVB/NJ mice were maintained in institutional vivarium and enrolled in experiments at 15-20 weeks of age. The subcutaneous allograft models were established by injecting 0.5 x 10^6^ cells (a generous gift from Steven Dubinett, UCLA) subcutaneously into a single flank. When tumors reached 80∼130 mm^3^, mice were randomized to saline control (PBS) or CRC2631/iSTORM treatment groups (4 intraperitoneal boluses of 5 x 10^7^ or 10^8^ CFU one every ∼3 day). Tumor dimensions length and width were measured bi-weekly using calipers, and tumor volume was calculated using the following formula 0.5 x L x W^2^.

Spatial Nanostring whole transcriptome profiling to assess the impact of bacteria treatment on the tumor microenvironment was performed by Bruker Spatial Biology (San Diego, California). Briefly, KPL tumors were harvested from mice treated either with PBS or CRC2631/iSTORM 4 days following treatment. Tumors were fixed in 10% neutral-buffered formalin (Epredia, Kalamazoo, MI, USA), transferred to 70% ethanol, and shipped to Bruker (San Diego CA) for sectioning, staining (anti-pan-cytokeratin/PanCK or anti-CD45 to detect tumor or immune cells, respectively), and imaging on a GeoMx DSP instrument across a minimum of 12 regions of interests (ROI ∼28mm^2^) selectively capturing either the tumor (PanCK+, CD45^-^) or the immune (PanCK^-^, CD45^+^) compartments per treatment condition. Comparative transcript expression analyses were performed using the GeoMx data analysis platform.

#### KPC GEMM tumor models in FVB/NJ immunocompetent mice

The genetically engineered pancreatic ductal adenocarcinoma (PDAC) model KPC (Kras^G12D/+^; Trp53^R172H/+^; Pdx1-Cre) on a C57BL/6 background was used to evaluate bacterial biodistribution and efficacy. Kras; Trp53; Tg(Pdx1-Cre/Esr1)#Dam/J mice (Jackson Laboratory, Bar Harbor, ME, USA; stock #032429) were bred, pups were weaned at day 21, and genotyped. Tumors were induced using tamoxifen prepared at 20 mg/mL in corn oil (Thermo Fisher Scientific; catalog# J63509.03). Mice 4-7 weeks of age received tamoxifen intraperitoneally at 75 mg/kg once daily for five consecutive days. MRI was used to verify tumor-positive status prior to downstream imaging or biodistribution studies. MRI was performed on a Bruker AVANCE III MRI system (Bruker BioSpin, Billerica, MA, USA). Mice were anesthetized with 3% isoflurane for induction and maintained at 1.5-2.5% isoflurane to sustain a respiratory rate of 26-35 breaths per minute. Respiratory rate was continuously monitored, and body temperature was maintained at 36-37°C using a heated imaging bed. Images were acquired using ParaVision 6 software (Bruker).

Following saline or bacterial treatment mice were monitored daily and pancreases were harvested 30-65 days post-treatment and weighed to assess impact of bacterial treatment on tumor growth.

### Toxicity study for iSTORM safety profile

Female C57BL/6J mice of 13-15 weeks of age were treated intraperitoneally with four doses (one every ∼3 days ) of saline control (PBS) or cultured iSTORM (5 x 10^7^ or 1 x 10^9^ CFU in 100 µL). Following the initial bolus, mouse weight, overall health status, and survival were monitored at least once daily. Peripheral blood was collected from the tail vein 12 days post-treatment initiation. 32 days after treatment initiation mice were euthanized by CO_2_ inhalation, and whole blood was collected via intracardiac puncture into K_3_EDTA- and sodium heparin-coated tubes (Sarstedt, catalog# 20.1288, NC, United States). 12-or 32-days blood samples were processed for cytokine quantification and blood chemistry, and immune cell profiling by flow cytometry. Livers and spleen were processed for histopathological analyses (see below).

Plasma was separated from whole blood collected in K₃EDTA-tubes (Sarstedt, catalog# 20.1288, NC, United States) at days 12 or 32 by centrifugation at 2000 rpm for 15 mins at room temperature (RT) and used in a customized Milliplex^®^ Multiplex Assay kit (Millipore, MA, United States). Results were acquired on a Magpix^®^ Multiplexing System (Luminex, TX, United States). Livers and spleen were processed for histopathological analyses.

#### Flow cytometry (safety study)

Blood cell pellet from K_3_EDTA-covered tubes at days 12 or 32 were first incubated with mouse F_c_ (CD16/32) blocker (BioLegend, catalog# 101320, CA, United States) for 10 mins at 4° C, and Fixable Viability Dye-BV421 (BioLegend, catalog# 423114) for 30 mins at RT. Cells were stained with the following antibody panel: CD3-AF488, CD4-AF700, CD8-PerCP-Cy5.5, CD19-PE/Dazzle^TM^ 549, Ly-6G-BV650, CD11b-BV750, F4/80-BV711, SiglecF-AF647, FceRI-PE-Cy**^®^**7, CD11c-PE. All the enlisted FACS antibodies have been purchased by BioLegend. After staining (30 mins at 4°C) and wash with FACS buffer (PBS supplemented with 3% FBS), blood cells were pelleted by centrifugation (200g for 6 mins at 4°C) and fixed with eBioscience^TM^ FoxP3 Transcription Factor Fixation/Permeabilization kit (Thermo Fisher Scientific, catalog# 00-5523-00) following manufacturer’s instructions. Stained samples were acquired on a 4L-spectral flow cytometer Aurora (Cytek Biosciences, CA, United States). After morphology and live cell gating, and exclusion of the doublets, gating system consisted in: CD4^+^ T cells (CD19^-^/CD3^+^/CD4^+^/CD8^-^), CD8+ T cells (CD19^-^/CD3^+^/CD4^+^/CD8^-^), B cells (CD19^+^/CD3^-^/CD11b^-^), Neutrophils (CD19^-^/CD3^-^/CD11b^hi^/Ly-6G^+^), Monocytes (CD19^-^/CD3^-^/CD11b^+^/F4/80^+^/Ly-6G^-^), Dendritic cells (DCs, CD19^-^/CD3^-^/CD11c^+^/FSC-A^low^/SSC-A^low^). Data were analyzed using FlowJo^TM^ software v10.0.7 (Tree Star, OR, United States).

#### Histopathology

Both kidneys and the right and left lateral liver lobes from controls or miced treated with iSTORM (5 x 10^7^ CFU) mice were harvested on day 32 post-treatment initiation and fixed overnight in 10% neutral buffered formalin (Epredia, MI, United States). The fixed tissues were then stored in 70% ethanol at 4°C, until submission to IDEXX BioAnalytics (Columbia MO, United States) for processing: paraffin embedding, H&E staining, and blinded analysis by a board-certified pathologist scoring necrosis, immune cell infiltrates, and extramedullary hematopoiesis. Tissues were trimmed according to RITA (Registry of Industrial Toxicology Animal-Data) protocols, and microscopic findings were graded for severity using a standard scoring system (0= no significant change, 1= minimal, 2= mild, 3= moderate, 4= severe) following INHAND (International Harmonization of Nomenclature and Diagnostic Criteria) standards.

### *In vivo* fluorescent imaging

All mice were fed 2019 Teklad global 19% protein extruded rodent diet (Inotiv, IN, United States) for 3 days before fluorescent imaging. This defined diet excludes alfalfa chlorophyll, minimizing feed-related autofluorescence in the gastrointestinal system^69^. CRC2631*^iRFP7^*^20-*cat*^ was reconstituted in 100 µL phosphate buffered saline (PBS) (Rocky Mountain Biologicals) and administered intravenously (i.v.) or in i.p. on day eight (female B6Panc02-H7 group) or day ten (male B6Panc02-H7 group) post-xenograft implantation. *In vivo* imaging was performed using a Xenogen IVIS 200 fluorescence system (Perkin-Elmer, MT, United States). Animals were handled in a biosafety cabinet where induction and maintenance of the anesthesia were delivered at 2.5-3% isoflurane in oxygen at approximately 1 L/min via a non-rebreathing inhalation system. When anesthesia level was appropriate (verified via toe pinch), animals were transferred from the induction box to the nose cones on the imaging platform. Images were acquired using the predefined filters for the fluorescent agent being used in the study. Animals were removed and immediately sacrificed to collect pancreas, Panc02-H7 xenograft tumors, liver, spleen, kidneys, lungs, and heart for *ex vivo* imaging, cryosection, and isolation of lymphocytes.

### Cryosection and microscopy

Cryosection was performed by using a cryostat HM525 NX Cryostat (Thermo Fisher Scientific). Harvested tumor or organ tissues were embedded in O.C.T. compound (Thermo Fisher Scientific, catalog# 23-730-571) and placed on dry ice immediately after. A 16 <u>μ</u>m-thick section of tissue was sliced and placed onto slides using a cryostat. 4% formalin solution was applied to fix these tissues. Nuclei were stained with DAPI (Thermo Fisher Scientific, #D1306) and imaged using a fluorescence microscope (Keyence, IL, United States). Tumors derived form syngeneic KPL mice treated with control or CRC2631 were fixed and stained with anti-salmonella antibodies (Novus Biologicals Cat# NB600-1087).

### Quantitative biodistribution analysis

C57BL/6 or B6Panc02-H7 orthotopic PDAC mice were injected intravenously (tail vein injections) with a single bolus of PBS or CRC2631*^iRFP7^*^20-*cat*.^ (2.5 x 10^7^ CFU). Mice were euthanized 96 hours post-injection. Whole blood, liver, primary pancreatic tumors and any discrete metastatic tumor masses were collected, weighed, and kept on ice. Whole blood samples were immediately diluted 1/10 in 25% glycerol and PBS and stored at - 80°C. Tissue samples were homogenized in 3 mL sterile PBS for 20 seconds on ice using a tissue homogenizer (TissueRaptor Qiagen, MD, United States) with sterile tips, mixed with 3 mL of sterile 50% glycerol, and stored at -80°C. All tissue samples were later thawed, passed through 40 μm-sterile filters (BD Biosciences, catalog# 22363547, NJ, United States), 20 µL dilutions spotted in triplicate on selective LB with 200 μg/mL thymine and 20 μg/mL chloramphenicol plates, incubated at 37°C and enumerated after 24 h following the Miles and Misra method^70^.

### Ex vivo imaging

All mice used in the biodistribution and toxicity studies were imaged on a Bruker AVANCE III MRI platform. This system has the capability of achieving a 50 μm-resolution for imaging tumor models. Mice were anesthetized using 3% isoflurane; anesthesia was maintained with 1.5-2.5% isoflurane to keep breathing rate at 26-35 bpm during which axial and coronal scans of the mouse body were performed. Images were taken using ParaVision 6 software (Bruker BioSpin Inc., MA, United States).

### Lyophilization of iSTORM

Lyophilization was performed by OPS Diagnostics (Lebanon, NJ, USA). Briefly, iSTORM cultures were expanded in LB broth supplemented with thiamine and ampicillin (24 h, 37°C, 160 rpm), harvested by centrifugation, and resuspended 1:1 in Microbial Freeze Drying Buffer (MFDB 500-06; OPS Diagnostics), a proprietary cryoprotectant formulation. Pre-lyophilization density was quantified by dilution-to-extinction, after which 500 µl aliquots were dispensed into sterile 5-ml amber serum vials, partially stoppered, and lyophilized using a shelf cycle consisting of freezing (-40°C, 2 h), primary drying (-15°C, 16 h, reduced pressure), and secondary drying (20°C, 2 h). Vials were sealed under vacuum, crimped, and stored at 4°C until use. Post-lyophilization viability was assessed after rehydration with 500 µl sterile water using both plate count CFU assays and dilution-to-extinction in 96-well plates. Lyophilized product was stored at 4°C until use and reconstituted in sterile PBS immediately prior to dosing.

### Anti-tumor Immune responses

#### KPC GEMM and orthotopic Panc02H7 mice

Immediately after harvesting, tumors were minced, followed by digestion in 0.04% collagenase IV (Gibco^TM^, catalog# 9001-12-1) and 0.02 mg/mL DNase I (Sigma, catalog# D5025) in GBSS (Sigma-Aldrich, catalog# G9779) with shaking at 240 rpm for 45 mins at 37°C. After digestion, samples were filtered using a 40 μm-strainer, then centrifuged. Red Blood cells (RBC) were lysed in RBC lysis Buffer (BD PharmLyse^TM^, catalog# 555899, NJ, United States) for 5 mins at 37°C, then centrifuged and resuspended in 0.04% BSA (in PBS) solution with fluorochrome-labeled antibodies. Isolation of spleen lymphocytes. Spleen was harvested from mouse and minced in DMEM (Gibco^TM^, catalog# 11965092, filtered with 40 <u>μ</u>m-nylon filter (Fisherbrand^TM^, catalog# 22-363-547) and centrifuged at low speed to pellet cells. Cell pellets were incubated with RBC lysis buffer (Thermo Fisher Scientific, catalog# 00-4333-57), then centrifuged to collect pelleted spleen lymphocytes. Antibody-stained cells were analyzed by FACS using a flow cytometer (BD Biosciences), and data analysis was performed with FlowJo^TM^ software (Tree Star; https://www.flowjo.com/). The entire cell population was gated with cell debris, then gated for live cells (i.e., 7AAD^-^) (BioLegend^®^, catalog# 420403). Next, CD4 and CD8 markers were used to distinguish CD4^+^ from CD8^+^ T cells and were gated for CD8-positive T cell activation or exhaustion markers (PD-1, LAG3, CD69…)

#### Subcutaneous KPC mice

Because tumor tissue availability was limited in the lyophilized iSTORM-treated cohorts, immune signatures were assessed in peripheral blood. PBMCs were isolated by density separation using Lymphopure, stained, and acquired on a Cytek Northern Lights cytometer. Data were analyzed in FlowJo.

Peripheral blood was collected from C57BL/6 mice under isoflurane anesthesia using heparinized capillary tubes. Approximately 200-300 µL of blood was obtained per mouse and immediately transferred into tubes containing 10 U/mL EDTA to prevent coagulation. Collected blood was diluted 1:1 with sterile phosphate-buffered saline (PBS, pH 7.4) and gently layered over an equal volume of Lymphopure™ (BioLegend, USA) in 15 mL conical tubes. Samples were centrifuged at 500 × g for 30 minutes at room temperature with the brake off to allow for optimal separation of blood components.

The mononuclear cell layer at the plasma-Lymphopure interface was carefully aspirated and transferred to a new tube. Cells were washed twice with PBS by centrifugation at 500 × g for 5 minutes to remove residual Lymphopure and platelets. The final pheripheral mononuclear cell (PBMC) pellet was resuspended in 100 µl of PBS (Thermo Fisher, USA) and 1% FCS (ThermoFisher, USA). Samples were stained with appropriate ratio (0.5 µl of live/dead stain: 10^6^ cells, 1µl of antibody: 10^6^ cells for the rest of antibodies) of fluorochrome-conjugated antibodies (table below).

Antibodies (Biolegend, USA) used for detection of tumors infiltrating leukocytes

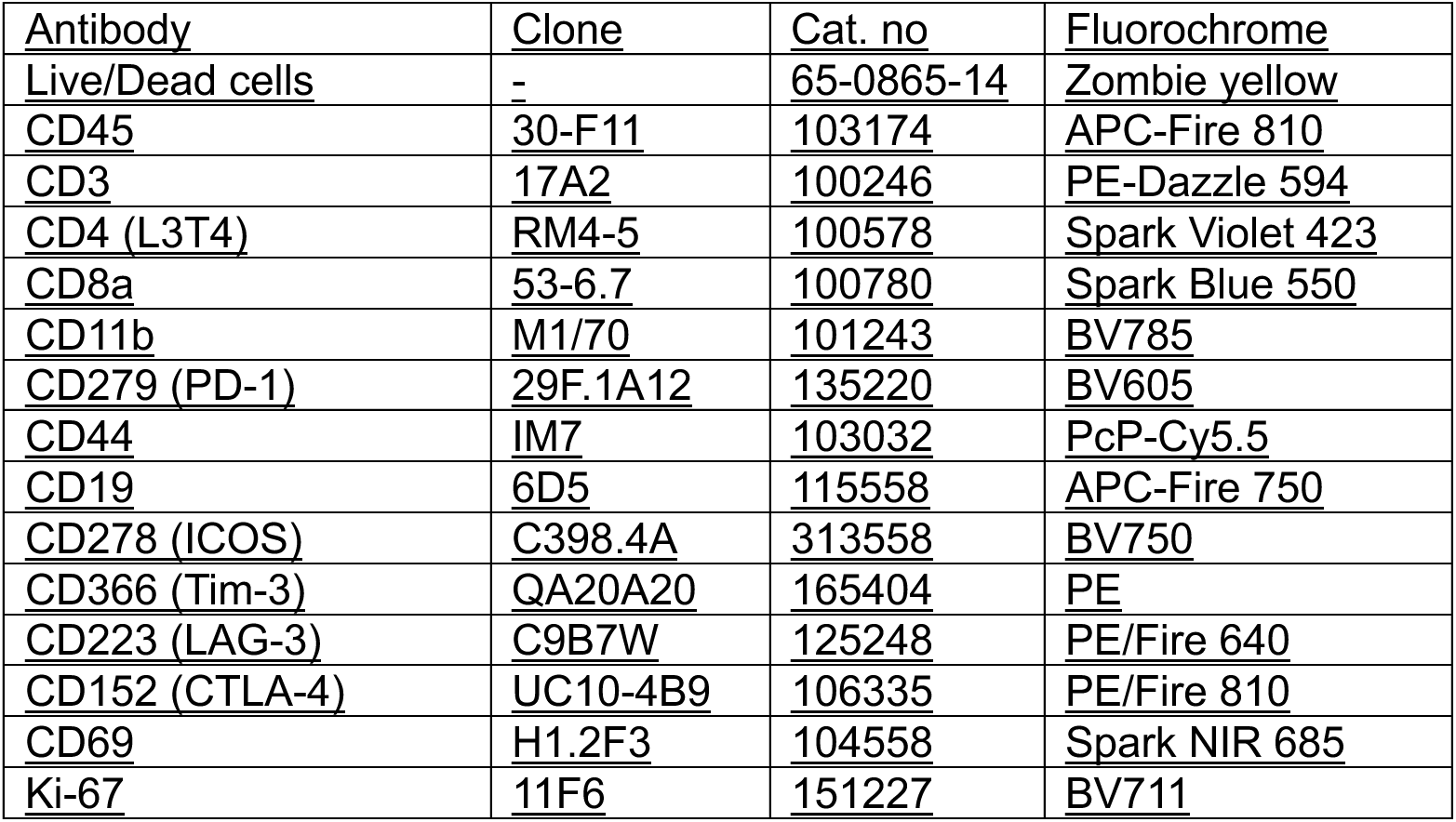

After 30min of incubation samples were washed 2 times with PBS (centrifuged at 500 g for 5 min at 25°C), the cell pellets resuspended in 100µl of PBS. Then the cells were fixed and analyzed using Cytek Northern Lights cytometer at Flow Cytometry Core of University of Arkansas for Medical Sciences. The results were analyzed using the FlowJo software v10.10 (BD Biosciences, San Jose, CA, USA).

#### Gating strategy

Cell size was used to define leukocyte populations before subset quantification. CD45 was used to identify immune cells. T cells were defined as CD45^+^CD3^+^. Within CD45^+^CD3^+^ cells, T helper cells were defined as CD4^+^CD8^-^ and cytotoxic T cells as CD4^-^CD8^+^. Activated T cells were identified using CD44, with additional markers including CD69, Ki-67, and ICOS. Co-inhibitory receptor phenotypes were assessed using PD-1, LAG-3, TIM-3, and CTLA-4. From CD45^+^CD3^−^ cells, CD19 and CD11b were used to stratify B cells and other lineages. Statistical analyses were performed in GraphPad Prism software (version 10.2.3; GraphPad Software, Inc., La Jolla, CA, USA). All results are presented as the mean ± SD. *, p < 0.05; **, p < 0.01; ***, p< 0.0005; **** p<0.00005 by a one-way ANOVA followed by the Dunnett’s multiple comparisons test.

### Lyophilized iSTORM tumor size control efficacy study

Subcutaneous (SQ) PDAC in syngeneic mice was generated by implanting 1.0 x 10^6^ KPC-luc cells into the right flanks of 6-8-week-old C57BL/6J mice. Tumor volumes in mice were monitored and measured using a digital caliper. Once tumors reached 80-120 mm^3^ mice were randomized to PBS or iSTORM treatments. Tumor size were measured twice weekly for 60 days.

### Additional statistical consideration

All statistical analyses (Log-rank Mantel Cox analysis of Kaplan-Meier curves, two-way repeated measure ANOVA for bacterial growth curves and weight loss graph Student’s *t* test analyses of CRC2631/iSTORM biodistribution, differences in tumor weight, immune profile, cytokine quantification and histopathologic score) were performed using GraphPad Prism software (v6.0 h and v10.6.1).

---

**Supplementary Figure 1. Dose-finding studies in the KPC genetic mouse model**

Comparative toxicological assessment of CRC2631 using mean percentage weight change of KPC GEMM mice treated with four 5 x 10^7^ CRC2631 dosage every three days either subcutaneous **(a)** or intraperitoneal (i.p.) **(b)** route of injection. Mean survival probabilities for these treatments are shown in panel **(c)**. The dose-dependent effects of CRC2631 (5 x 10^7^, 2 x 10^8^, and 5 x 10^9^ CFU) with PBS as a control were evaluated as well tumor-bearing KPC mice. N value represents the number of mice. Mouse weights were recorded daily.

**Supplementary Figure 2. CRC2631 and iSTORM selectively colonize lung tumors**

**(a-b)** CRC2631 colonizes tumor cores. 15-20 weeks-old FVB/NJ mice were subcutaneously inoculated in the flank with 0.5 x 10^6^ KPL (Kras^G12D/+^; STK11^-/-^; ROSA26*^LSL-Luc^*) murine cells. Tumors-positive (60 mm^3^) FVB/NJ mice were treated intraperitoneally with a single dose CRC2631 or PBS vehicle control. Tumor allografts were harvested 4 days post-treatment, fixed and paraffin embedded, sectioned, and stained with anti-Salmonella antibodies (Novus Biologicals, #NB600-1087, CO, United States). Deep tissue sections show tumor-specific CRC2631 positivity.

**(c-e)** Similar to its demonstrated avidity for pancreatic tumors, iSTORM targets lung tumors in the subcutaneous KPL model. When tumors reached ∼130 mm^3^, mice were randomized to saline control (PBS) or iSTORM treatment groups (*N*= 6/group, 4 boluses of 5 x 10^7^ CFU, or 10^8^ one every ∼3 day/full dose). Tumors (**c**) and indicated visceral organs (**d**, **e**) were harvested 1 day following full treatment and subjected to ex-vivo imaging.

**Supplementary Figure 3. CRC2631 Induces Immune recruitment axis and antigen presentation signatures that are accompanied by host apoptosis signals in lung tumors**

Subcutaneous KPL lung tumors from mice treated intraperitoneally with PBS or CRC2631 were harvested 4 days after treatment and processed for spatial transcriptomic (GeoMx).

(**a-e**) Chemokine & adhesion switch. Cx3cl1, Cxcl10, Cxcl9, Icam1, Vcam1 rise 3- to 14-fold, signaling CXCR3-directed effector recruitment.

(**f-l**) Antigen-presentation stimulation. up-regulation of H2-K1, H2-D1, B2m, Tap1, Erap1, Psmb8/9 confirms heightened MHC-I processing and display capacity.

(**n-p, s, t**) IFN/TLR signaling Pathway heat-maps and STAT/Irf transcripts document robust TLR4-to-NF-κB and type-I/II-IFN cascades triggered by bacterial PAMPs and DAMPs.

(**m, r**) Concurrent elevation of Cd86 and Pd-l1 (Cd274) indicates productive APC activation but emergent adaptive resistance.

(**q, u**) Inflammasome / death signature. Nlrc4 and cell death genes (Casp3, Gsdmd, Ripk1, etc.) validate inflammasome engagement and on-target tumor cell killing.

**Supplementary Figure 4 iSTORM-induced anti-tumor immune signatures are superior to CRC2631**

(**a-j**) Spatial omics transcriptomic data showing the effect of treatment on the expression of the indicated genes in the tumor compartment (PanCK^+^) or in TME. FVB mice harboring syngeneic KPL flank tumors were treated with CRC2631 or iSTORM (5 x 10^7^ CFU, in i.p.). Tumors were harvested, fixed, and processed for spatial whole transcriptomic analyses. Up to 24 regions of interest from independent animals/treatment conditions were analyzed. (**a**) Compared to CRC2631 control, iSTORM reduced CMTM6 mRNA levels in the tumor compartment. (**b**) High levels of CD45 expression were noted in the compartment abutting tumors. (**c and d**) antigen cross presentation was among the top pathways upregulated following treatment with CRC2631 or iSTORM. These signatures are more robust in iSTORM compared to CRC2631-treated mice.

(**e, h-l**) Cell profiling deconvolution shows treatments convert the TME into an immunologically active state (**e**) characterized by elevated proportions of, dendritic cells/DC, natural killers/NK, natural killer T-cells/NKT, macrophages, and pro-inflammatory responses (**h-l**).

(**f**) Volcano plot showing comparative expression of CRC2631/iSTORM induced genes. iSTORM triggers a dramatic increase in the expression of tumor cytotoxic granzymes compared to CRC2631, highlighting iSTORM superior immune-stimulating capabilities. Comparative expression of granzyme-e (Gzm-e) is presented in “**g**” as an example.

(**h-k**) Spatial transcriptomic analysis in the tumor-associated immune compartment reveals iSTORM-driven immune reprograming transcriptional signatures. Nfkb1 (**h**) was elevated in CRC263/iSTORM-treated tumors, indicating enhanced NF-κB-dependent innate activation. Stat1,

(**i**) and the macrophage-associated interferon regulators Irf5 (**j**) and Irf8 (**k**) were upregulated, consistent with type I interferon signaling and pro-inflammatory macrophage polarization.

(**l**) Violin plots showing the effect of iSTORM treatment on tumor growth. 15-20 weeks-old FVB/NJ mice harboring subcutaneous KPL tumors (130 mm^3^) were treated i.p either with saline control (PBS) or iSTORM (*N*= ∼6/group, 3 boluses of 10^8^ CFU, one every ∼3 day). Caliper tumor measurements were used to derive on-treatment percentage tumor size changes from baseline up to 1 day after the last treatment. Each dot represents an individual mouse; dashed lines indicate the median. PBS-treated tumors exhibit robust progression, whereas iSTORM treatment shifts the response distribution toward tumor stasis and regression. Statistical significance was assessed using a two-tailed Mann-Whitney U test (*p* = 0.028). The horizontal dashed line denotes 0% change (no growth).

**Supplementary Figure 5. iSTORM shows favorable safety profile in excretive organs**

Graphs show histopathologic scoring for immune cell (IC) infiltrates, extramedullary hemopoiesis (EMH) and necrosis assessed in kidneys and livers harvested from C57BL/6 mice treated either with from PBS or 5 x 10^7^ CFU (*N*=3/group). A statistically significant increase of IC infiltrates was observed in the kidney but limited to the perirenal adipose tissue. There was no evidence of IC infiltrates in the parenchyma of the kidneys. Error bars show standard deviation and p-values are derived from Student’s t-test.

**Supplementary Figure 6. Safety and tolerability of iSTORM in aged mice.**

Age-sensitized (58-85-weeks old males and female) FVB/nJ mice received four intravenous doses of saline control (PBS) (yellow, *N= 3*) or lyophilized iSTORM (iSTORM-L) on days 0, 3, 6, and 10 (green arrowheads) at either 5 x 10^7^ CFU (purple, *N= 3*) or 5 x 10^8^ CFU (light blue, *N= 4*).

**(a)** Body weight was recorded daily from the start of the treatment and expressed as percent change relative to baseline (day 0), which did not reveal any significant changes across the arms of the cohort. (**b**) Survival was monitored for 32 days. Dose-dependent lethality was observed in the high dose (5 x 10^8^ CFU) cohort. Error bars show standard deviation and p-values are derived from two-way repeated measure ANOVA **(a)** and Log-Rank test **(b)**.

**(c)** Blood from the animal cohort above was collected on day 12 after treatment (red arrow) to assess immune toxicity. iSTORM-L didn’t drastically alter hematic analytes. Error bars show standard deviation and p-values are derived from Student’s t-test.

**Supplementary Figure 7. Comparative growth kinetic analysis of standard versus lyophilized iSTORM**

The growth curves highlight a bigger growing trend (*p<*0.05) in the cultured strain until 16 hours of incubation in respect to the lyophilized (green line) strain. This justifies the inclusion of a higher dose (1 x 10^8^) of lyophilized iSTORM for anti-tumor treatment in addition to the 5 x 10^7^ CFU/mouse dose. Each condition was assessed in biological triplicates. Error bars show standard deviation, and p-values are derived from two-way repeated measure ANOVA test.

**Supplementary Figure 8. Potential role of target cell mannose-rich glycans in iSTORM tumor avidity**

(**a**) Diagram describing the high-throughput glycan-binding array performed against CRC2631 strain. (**b**) Results of the glycan-binding screening showed that the attenuated bacterial strain binds with more affinity to mannose-rich N-glycans, including transferrin and transferrin receptor-linked structures. (**c, d**) Western blot analysis of the Transferrin Receptor (TfR) in hTERT-HPNE and CFPAC-1 cells. TfR expression was considerably higher in PDAC cell lines (CFPAC and PANC-1) in CFPAC compared to the control (HPNE). (**d**) PANC and HPNE were treated with PNGase and the protein size shift was detected by Western blot. (**e**) PANC-1, KPCY and hTERT-HPNE were treated with aTfR and then they were cocultured with CRC2631-iRFP at 10:1 MOI. Cell viability was measured using Trypan blue staining. Total number of viable cells were used to calculate the percentage of reduction in cell killing.

