## Supplementary Figures for "CMTM6-Silencing Microbial Immunotherapy Reprograms PDAC Tumors and Restores T-cell Function"

### Supplementary Figure 1

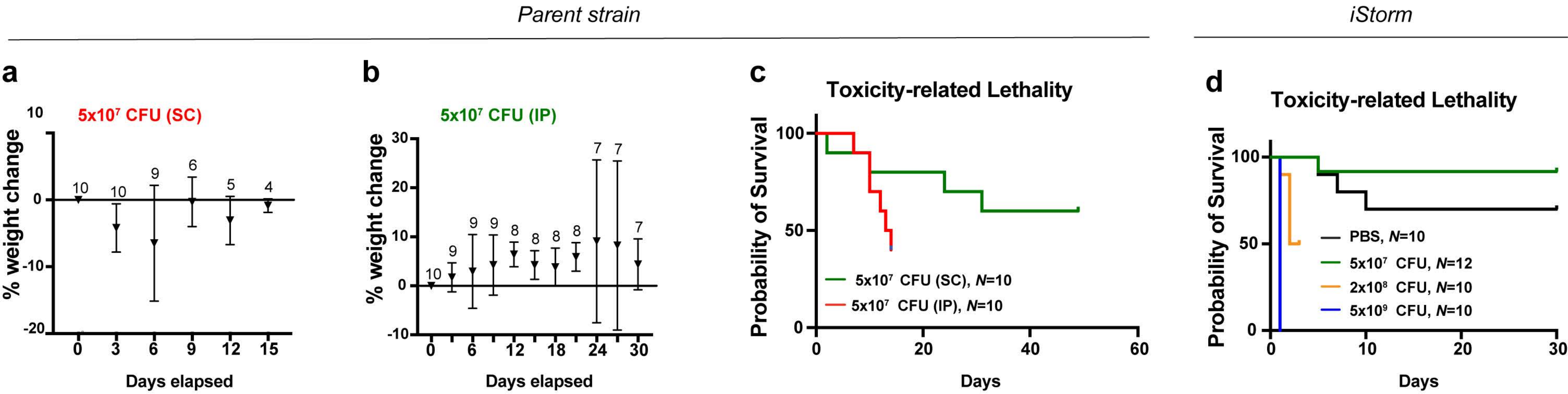

Supplementary Figure 2

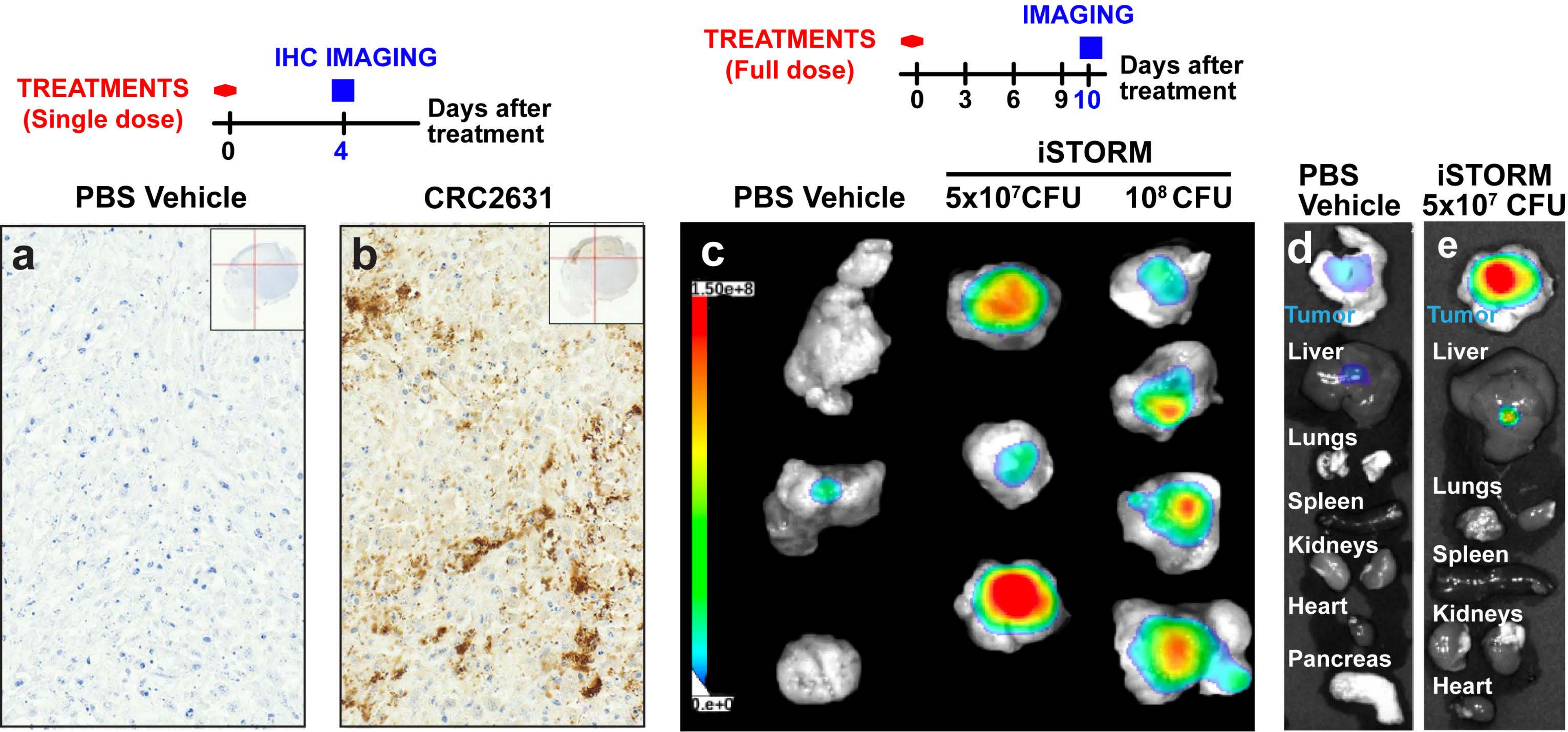

### Supplementary Figure 3

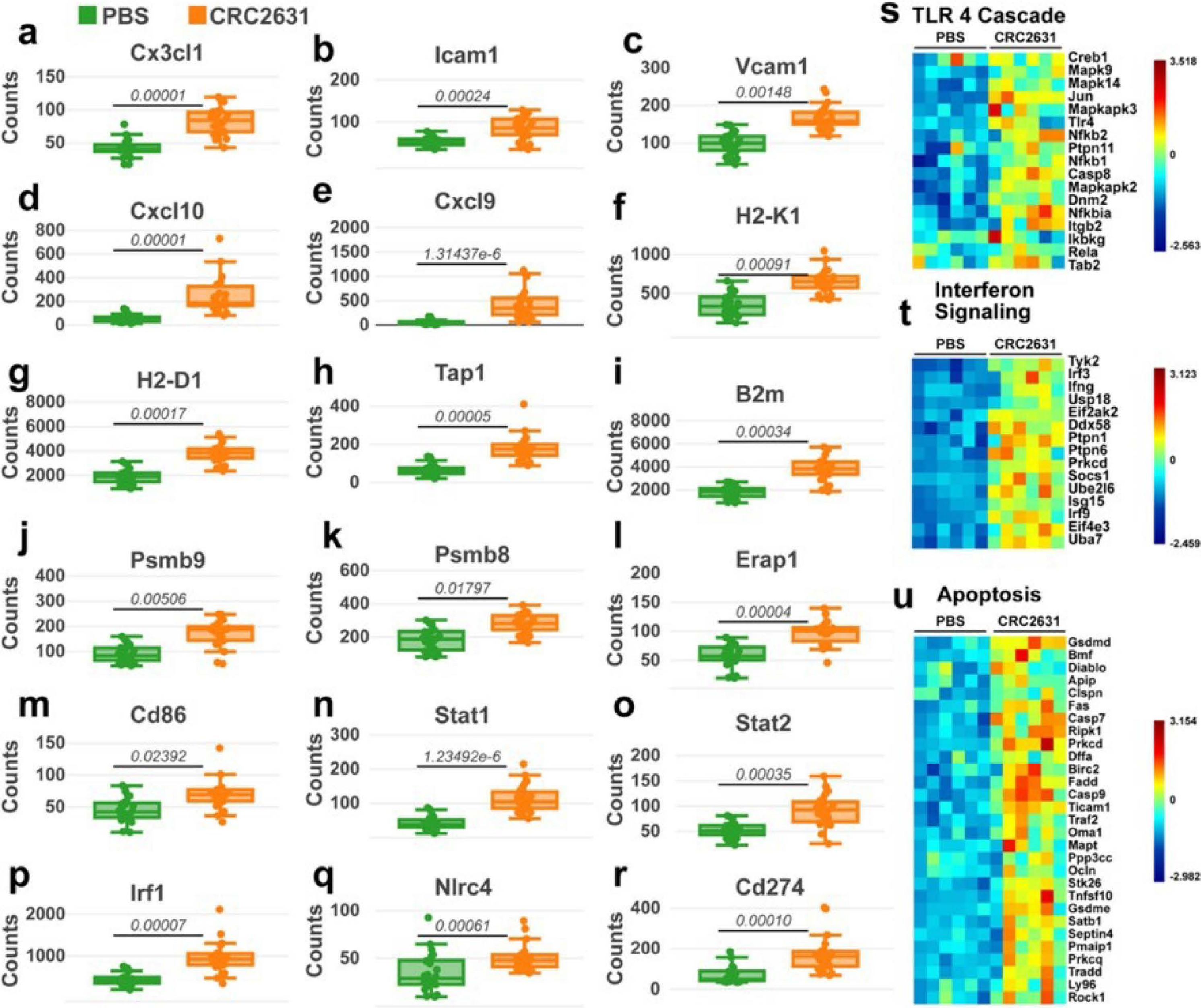

### Supplementary Figure 4

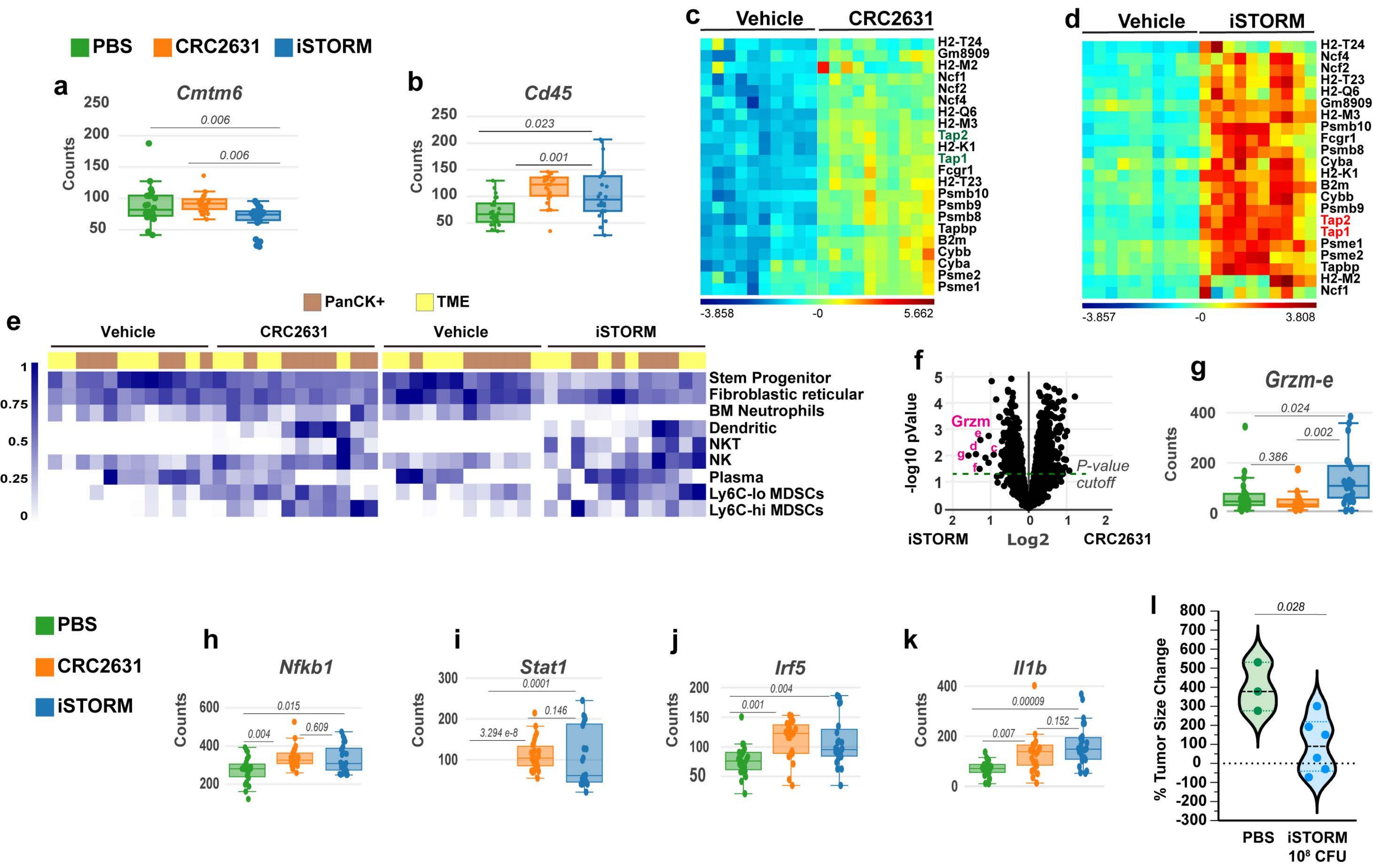

#### Supplementary Figure 5

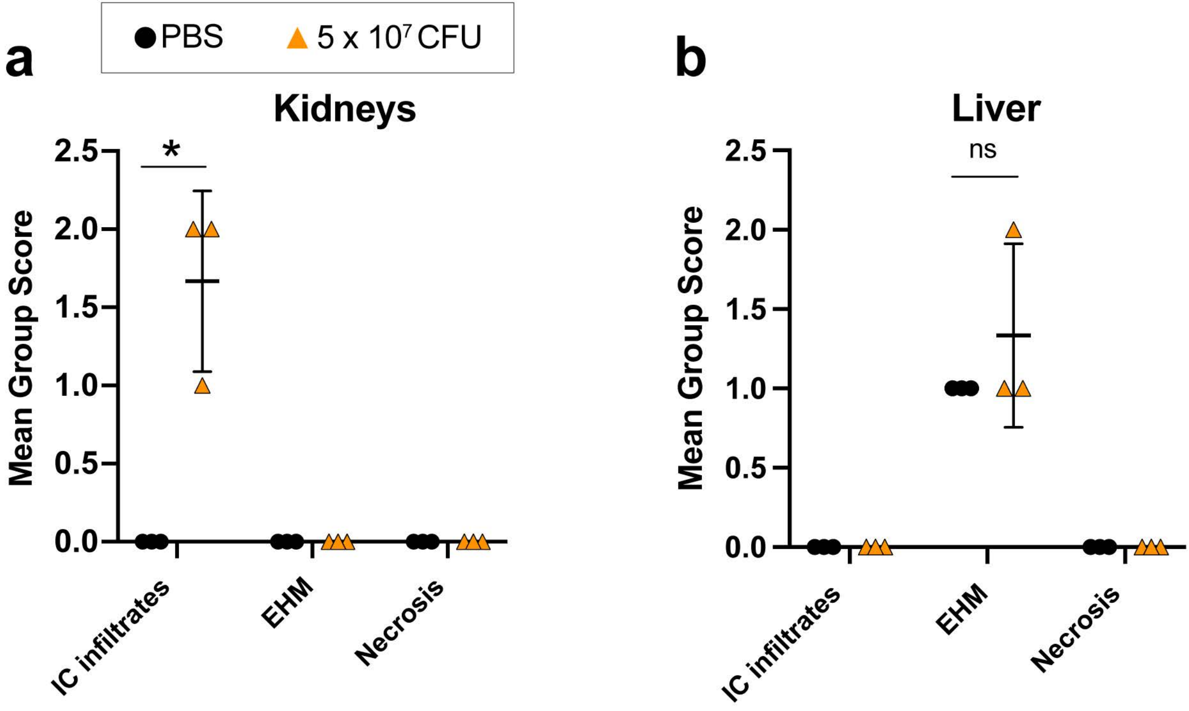

### Supplementary Figure 6

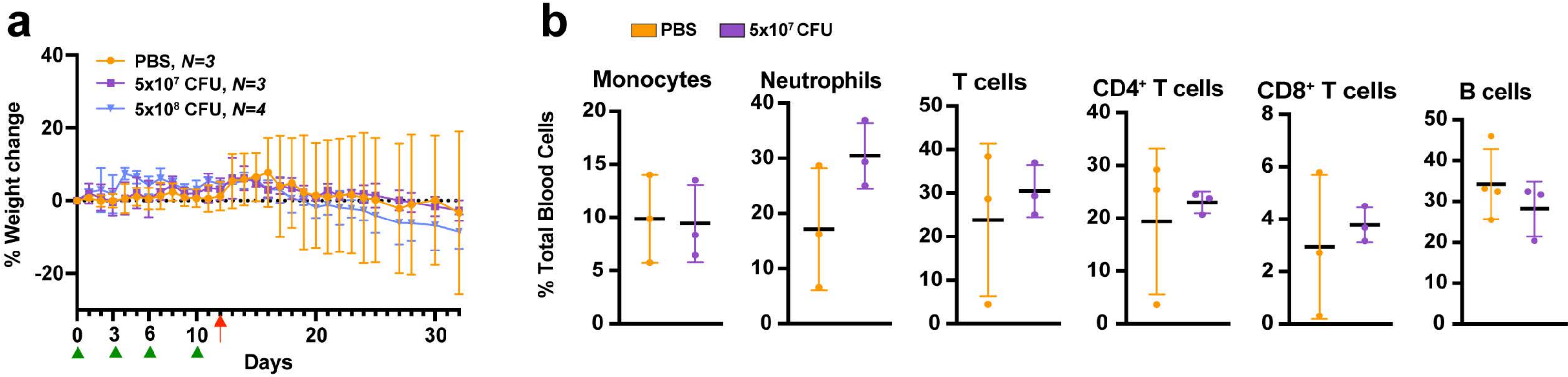

**Supplementary Figure 7**

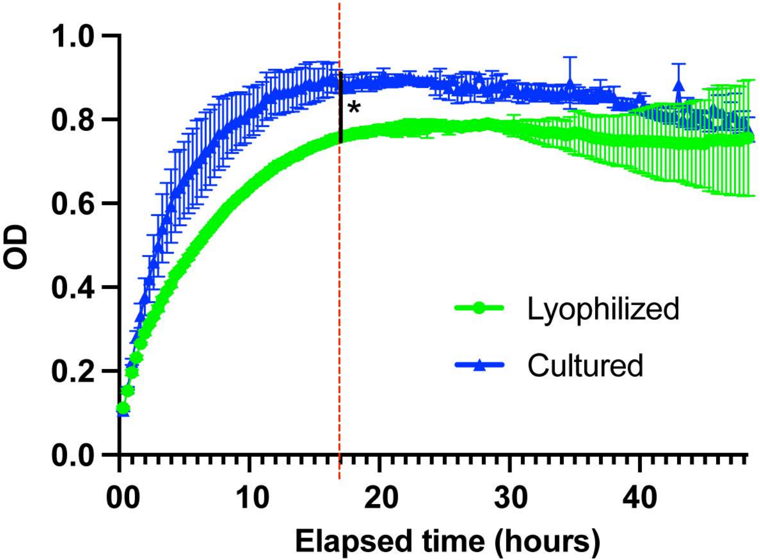
